# High-speed mapping of whole-mouse peripheral nerves at subcellular resolution

**DOI:** 10.1101/2025.01.22.632569

**Authors:** Mei-Yu Shi, Yuchen Yao, Miao Wang, Qi Yang, Lufeng Ding, Rui Li, Yuanyuan Li, Haimeng Huang, Chao-Yu Yang, Zhao Zhou, Zhenxiang Zhu, Pengjie Wen, Fangling Dai, Xiaohui Zeng, Ke-Ming Zhang, Yuhong Guo, Zi-An Sun, Huanhuan Xia, Zhenhua Ren, Yusuf Ozgur Cakmak, Ming Zhang, Fuqiang Xu, Lei Qu, Qingyuan Zhu, Pak-Ming Lau, Cheng Xu, Guo-Qiang Bi

**Affiliations:** Hefei National Research Center for Physical Sciences at the Microscale, University of Science and Technology of China, Hefei 230026, Anhui, China; Anhui Province Key Laboratory of Biomedical Imaging and Intelligent Processing, Institute of Artificial Intelligence, Hefei Comprehensive National Science Center, Hefei 230088, Anhui, China; Division of Life Sciences and Medicine, CAS Key Laboratory of Brain Function and Disease, University of Science and Technology of China, Hefei 230026, Anhui, China; Institute of Advanced Technology, University of Science and Technology of China, Hefei 230026, Anhui, China; The Brain Cognition and Brain Disease Institute, Shenzhen-Hong Kong Institute of Brain Science-Shenzhen Fundamental Research Institutions, Shenzhen Institutes of Advanced Technology, Chinese Academy of Sciences, Shenzhen 518055, Guangdong, China; Shenzhen Bineogen Technology Co., Ltd, Shenzhen 518107, Guangdong, China; Ministry of Education Key Laboratory of Intelligent Computation and Signal Processing, Information Materials and Intelligent Sensing Laboratory of Anhui Province, School of Electronics and Information Engineering, Anhui University, Hefei 230601, Anhui, China; SEU-ALLEN Joint Center, Institute for Brain and Intelligence, Southeast University, Nanjing 211189, Jiangsu, China; Anhui Communications Vocational and Technical College, Hefei 230051, Anhui, China; Shenzhen Key Laboratory of Viral Vectors for Biomedicine, NMPA Key Laboratory for Research and Evaluation of Viral Vector Technology in Cell and Gene Therapy Medicinal Products, Shenzhen Institutes of Advanced Technology, Chinese Academy of Sciences, Shenzhen 518055, Guangdong, China; State Key Laboratory of Magnetic Resonance and Atomic and Molecular Physics, Key Laboratory of Magnetic Resonance in Biological Systems, Wuhan Center for Magnetic Resonance, Innovation Academy for Precision Measurement Science and Technology, Chinese Academy of Sciences, Wuhan 430071, Hubei, China; University of Chinese Academy of Sciences, Beijing 100049, China; Department of Anatomy, University of Otago, Dunedin 9054, New Zealand; School of Medicine, The Chinese University of Hong Kong (Shenzhen), Shenzhen 518172, Guangdong, China; Department of Anatomy, School of Basic Medicine, Anhui Medical University, Hefei 230032, Anhui, China

**Keywords:** whole-body imaging, tissue clearing, VISoR, peripheral nervous system, spinal nerve, sympathetic nerve, vagus nerve

## Abstract

In contrast to the rapid advancements in mesoscale connectomic mapping of the mammalian brain, similar mapping the peripheral nervous system has remained challenging due to the body’s size and complexity. Here, we present a high-speed blockface volumetric imaging system with an optimized workflow of whole-body clearing, capable of imaging the entire adult mouse at micrometer resolution within 40 hours. Three-dimensional reconstruction of individual spinal fibers in Thy1-EGFP mice reveals distinct morphological features of sensory and motor projections along the ventral and dorsal rami. Immunostaining facilitates body-wide mapping of sympathetic nerves and their branches, highlighting their perivascular patterns in limb muscles, bones, and most visceral organs. Viral tracing elucidates the fine architecture of vagus nerves and individual vagal fibers, revealing unexpected projection routes to various organs. Our approach offers an effective means to achieve a holistic understanding of cellular-level interactions among different systems that underlie body physiology and disease.

**Highlights:**

- Imaging of whole adult mouse at uniform subcellular resolution within 40 hours
- Reconstruction of 191 thoracic spinal neuron projections for Thy1-EGFP mice
- Mapping of immunolabeled sympathetic projections and their perivascular patterns
- Resolving virus-labeled architecture of vagus nerves and complex single-fiber routes

## Introduction

The nervous system, consisting of the central nervous system (CNS) and peripheral nervous system (PNS), is an intricate network of interconnected neurons that spans the entire body, coordinating complex physiological and psychological activities.^1–5^ Visualizing and mapping the anatomical structures of this complex system is essential for gaining a mechanistic understanding of its functions and associated diseases.^6,7^ Over the past decade, significant progress has been made in mesoscale connectomic mapping of the CNS at subcellular resolution across the whole brain, largely due to advances in three-dimensional (3D) optical microscopy.^8–16^ However, similar analyses for the PNS have remained challenging.^6,17,18^ Unlike the relatively compact and homogeneous brain, the mammalian body is much larger and highly heterogeneous, containing irregular structures and diverse tissue types.^7,19^ As a result, even with state-of-the-art imaging techniques, resolving the long, complex, and often intermingled nerve routes of the PNS at the whole-body scale remains extremely difficult.^6,7^ Consequently, our current architectural understanding of the PNS is still largely based on gross anatomical observations, which provide a general but imprecise overview of its routing, branching, and connecting patterns.^20–22^

Recent technical advances have attempted to visualize peripheral nerves alongside other body structures by utilizing whole-body clearing techniques with improved solvent- and aqueous-based methods.^23–28^ These include DISCO series especially wildDISCO that achieves whole body visualization of immunolabeled mice,^23,28^ and HYBRiD and TESOS that achieve whole body visualization of fluorescent protein-expressing transgenic mice.^24,26^ Combined with conventional light-sheet microscopy, these clearing approaches enable coarse visualization of the overall nerve architecture and major branches of the PNS, although fine details, such as thin nerve branches and single axonal fibers, could not be uniformly resolved throughout the whole mouse body. Using confocal microscopy in a blockface imaging mode, Yi *et al*. achieved sub-cellular resolution to trace individual axons from the forepaw to the spinal cord in TESOS-cleared mouse body parts.^24^ Yet, the limited speed of confocal microscopy and block-milling hinders its scalability for high-resolution imaging of larger samples like the whole adult mouse body. Therefore, a high-speed method capable of efficiently imaging the entire adult mouse at uniformly high resolution is needed for mesoscale connectomic mapping of both the CNS and PNS. Ideally, this method should be compatible with commonly used fluorescence labeling techniques, such as immunostaining and viral labeling, to facilitate the visualization of neurons and their interactions with other tissues and cell types throughout the body.

A key technical challenge in optical imaging of large block samples, such as the mouse body, is the need for long working distance lenses to image deep within the sample. This requirement results in a lower numerical aperture, and consequently, lower resolution. Additionally, imperfect refractive index matching, due to tissue heterogeneity, causes aberrations that accumulate exponentially as light penetrates deeper tissues, further degrading image quality. Another major obstacle is the time required for data acquisition, which becomes a bottleneck in high-resolution imaging of large volumes.^12,24^ In recent years, we developed a novel light-sheet imaging method called volumetric imaging with synchronized on-the-fly scan and readout (VISoR), which enables ultrahigh-speed 3D imaging of cleared brain sections.^11,29^ Through computational reconstruction of the entire volume from serial section images, VISoR has proven to be an efficient tool for mesoscale mapping of mouse and monkey brains at micron resolution, allowing brain-wide tracing of individual axons.^11,29^ In theory, this approach could be adapted for high-resolution whole-body imaging. However, sectioning the mouse body presents significant challenges. The highly heterogeneous body structure, particularly the extensive natural cavities, often leads to deformation and damage of body slices prior to VISoR imaging.

To overcome these challenges, we developed an integrative approach comprising three key components: (1) an efficient procedure for uniform whole-body clearing and versatile labeling, (2) a blockface-VISoR system for serial sectioning with ultrahigh-speed 3D imaging of the cut surface, and (3) an algorithm and software for automated volume reconstruction from the serial 3D image set (Figure 1A). This method enables uniform subcellular imaging of an entire adult mouse within 40 hours. By applying this pipeline alongside transgenic labeling, whole-body immunostaining, or adeno-associated virus (AAV) labeling, we successfully imaged a few dozens of adult mice at micron resolution. Our approach revealed the complex architecture of the PNS, including spinal somatic motor and sensory nerves, the visceral sympathetic and vagus nerves, and their interactions with various none-neural tissues and organs throughout the body.

**Figure 1.**
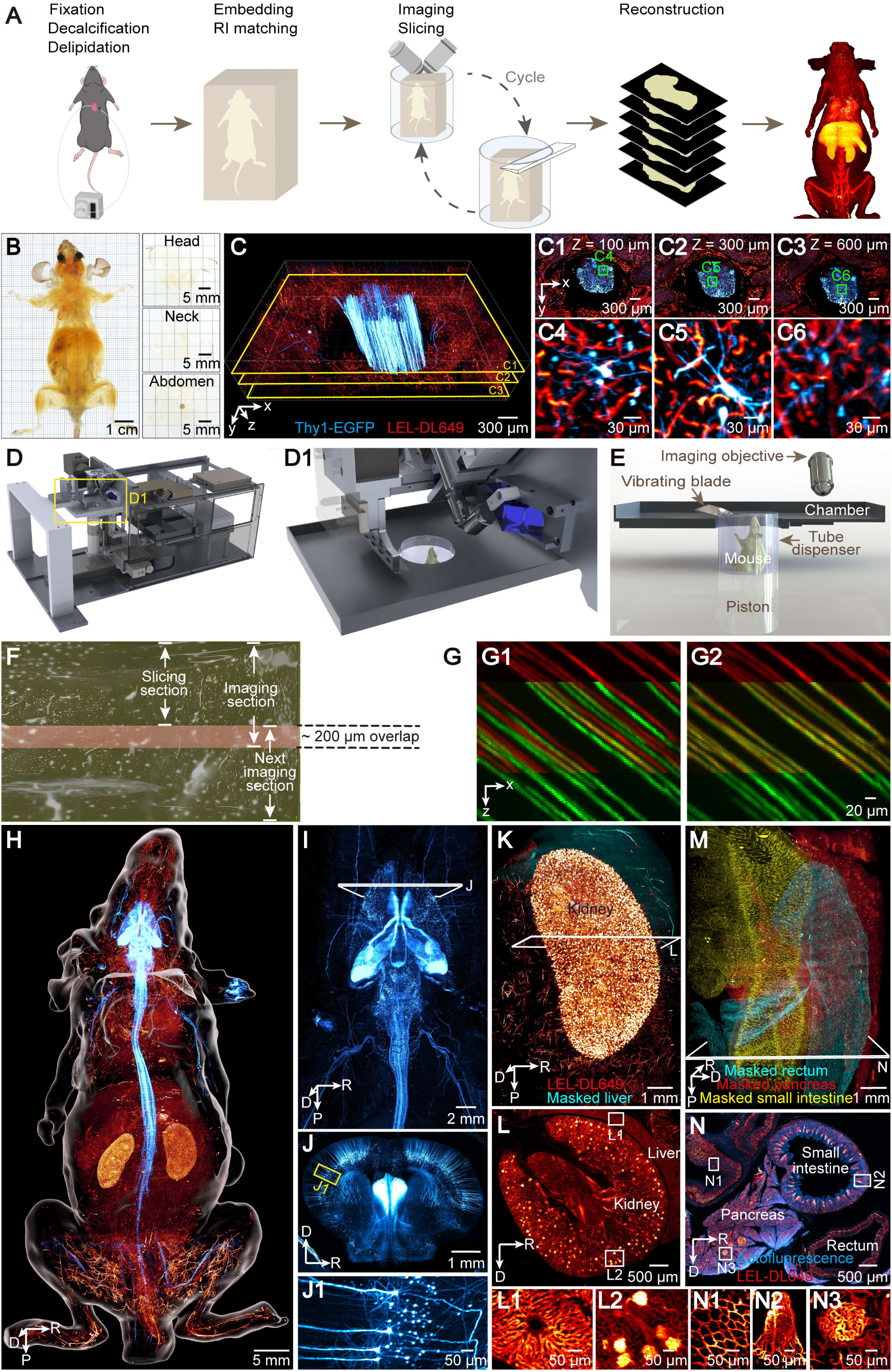
ARCHmap-blockface-VISoR approach for high-throughput mapping of whole mouse body at micron resolution. (A) ARCHmap-blockface-VISoR pipeline for whole-mouse imaging. (B) Bright-field view of a cleared 2-month-old vGAT-Cre;Ai14 mouse and 400-μm-thick sections collected during blockface imaging. (C) 3D view of a ∼600-μm-thick thoracic volume imaged from a blockface imaging section. (C1-C6) Representative high-resolution images showing maximum intensity projection of 20-μm-thick stacks in (C) at depths of 100 μm (C1, zoomed-in view in C4), 300 μm (C2, zoomed-in view in C5), and 600 μm (C3, zoomed-in view in C6). (D, D1) Blockface-VISoR setup. (D1) Magnification of the imaging region in (D). (E) Cross section of imaging and sectioning modules of the blockface-VISoR setup. (F) Schematic showing overlapped imaging data (pink) of ∼200-µm thickness between 2 contiguous imaging sections. (G) Comparison of 3D intersection alignment by natural coordinates (G1) and a custom 3D reconstruction method (G2). Images of vessel fluorescence are the maximum intensity y-projection of 20-µm-thick sections in the upper section (red) and the lower section (green). (H) 3D view of nerve and vasculature in a whole adult Thy1-EGFP mouse. Red hot (black-red-yellow-light yellow spectrum), LEL-DL649; cyan hot (black-blue-cyan-white spectrum), Thy1-EGFP; white surface, whole mouse. (I) 3D view of the nervous system in mouse head and thorax. (J) Maximum intensity projection of a 400-μm-thick stack in (I). (J1) Enlargement of the selected region in (J) showing the cortical neurons and fibers. (K) 3D view of vasculature in the kidney. (L) Maximum intensity projection of 20-μm coronal sections in (K) showing vasculature in the kidney and liver. (L1, L2) Magnified hepatic sinusoid (L1) and renal corpuscles (L2) in (L). (M) 3D view of vasculature in visceral organs segmented by pseudo colors. Cyan, large intestine; yellow, small intestine; red, pancreas. (N) Maximum intensity projection of 50-μm coronal sections in (M). (N1-N3) Magnified vasculatures in the submucosal layer of the large intestine (N1), small intestinal villi (N2), and islets of Langerhans in the pancreas (N3) in (N). Orientation of images: R: right; P: posterior; D: dorsal.

## Results

### ARCHmap: an aqueous-based whole mouse clearing pipeline for 3D blockface imaging

High-quality sample preparation is critical for large-volume imaging. Instead of relying on solvent-based whole-body clearing methods used in prior studies,^23,30–34^ we opted for an aqueous-based approach to better preserve signals from natively expressed fluorescent proteins.^27,35–38^ After evaluating various established techniques on extracted organs,^39–47^ we selected the CUBIC-L method and developed a modified protocol (hereafter referred to as CUBIC-LH, see Methods) that achieves excellent tissue clearing and fluorescent signal preservation with minimal tissue size change (Figure S1A-S1E).

We employed continuous active intracardiac perfusion for whole-body fixation, decalcification, and clearing with CUBIC-LH. This method, compared to traditional passive postfixation at low temperatures, resulted in significantly better permeation, especially in the brain and spinal cord (Figure S1F-H). To address sample deformation during sectioning, we developed an optimized hydrogel embedding solution, containing hydrogel monomer solution (HMS), bovine serum albumin (BSA), and iohexol. This recipe greatly improved the mechanical strength of the cleared mouse body (Figure S2A). Finally, by using our previously developed PuClear refractive index (RI) matching solution,^29^ excellent optical transparency was achieved across the entire mouse body, enabling clear visualization of structures at least 600 μm below the body surface (Figure 1B-C and S2B).

In summary, we established a highly efficient aqueous-based protocol—**A**ctive Clea**R**ing with **C**UBIC-LH, **H**ydrogel-embedding, and RI-**ma**tching with **P**uClear (ARCHmap)—that provides homogeneous clearing of the entire mouse body.

### Blockface-VISoR imaging and mesoscale 3D reconstruction of the whole mouse body

To achieve uniform high-resolution imaging of the cleared mouse body, we developed the blockface-VISoR system, which integrates an imaging module with a precision sectioning module (Figure 1D). The imaging module features a custom-built fluorescence microscope utilizing our previously developed ultrahigh-speed VISoR technology.^11,29^ This system includes a laser light source, a galvanometer, a V-shaped pair of objective lenses, a high-speed scientific CMOS camera, and a precision x-y translation stage. The sectioning module is equipped with a custom-designed vibratome positioned next to the imaging head and a tube dispenser mounted on the microscope’s translation stage. The sample is loaded into the tube dispenser, which advances the sample at micron-precision increments (Figure 1D-1E and Video S1A). During operation, the blockface-VISoR system images only the tissue hundreds of microns below the freshly cut surface (blockface) of the cleared sample, allowing for consistently high-resolution imaging across the entire sample (Figure 1C).

High-throughput whole-body imaging is performed through automated cycles of serial sectioning and blockface imaging (Figure 1A, Video S1). At the start of each cycle, the top of the transparent mouse body, held within the tube dispenser, is pushed out for the vibratome to create a flat blockface. The microscope then captures 3D image sets in columns of oblique views,^11^ typically extending 600 μm below the blockface, at a rate of 200 fps while the translation stage moves the sample at a steady 0.7 mm/s (Video S1B). After imaging, the sample returns to the sectioning zone, where it advances by 400 μm for the vibratome to make the next cut, then moves back to the imaging zone for the next cycle (Figure 1E, Video S1). The 200-μm overlap between imaged sections ensures continuous coverage and is used to facilitate alignment during reconstruction (Figure 1F). Each cycle takes approximately 5-15 minutes for imaging and 3 minutes for sectioning. In total, a complete mouse body (∼ 4 × 2 × 8 cm³) can be imaged at a voxel resolution of 1 × 1 × 2.5 μm³ in about 40 hours (∼200 cycles), generating approximately 70 terabytes of data per fluorescence channel.

To reconstruct the entire mouse body, we adapted our previously developed software framework,^29^ and implemented a newly developed method based on the minimization of the normalized cross-correlation (NCC) between overlapping sub-volumes to improve inter-section stitching (see Methods). This method generates non-rigid deformation fields from NCC-based displacement parameters, which are then used in a global optimization process. This approach enables precise 3D alignment between adjacent imaged sections (Figure 1G, S3). As a demonstration, we reconstructed the whole body of an adult Thy1-EGFP mouse with vasculature labeled using tomato lectin conjugated with DyLight 649 (LEL-DL649) (Figure 1H, Video S2). The consistently high resolution across the dataset is evident in the zoomed-in views of various organs. These include individual neurons in the brain (Figure 1I-1J), radial vasculature in the hepatic sinusoids of the liver (Figure 1K-1L, 1L1), glomeruli in the renal corpuscles of the kidney (Figure 1K-1L, 1L2), capillary networks in the mucosa and submucosa layers of the intestine (Figure 1M-1N, 1N1-1N2), and Langerhans’ islets within the exocrine parenchyma of the pancreas (Figure 1M-1N, 1N3), all from the same 3D dataset.

### Visualization of cranial nerves in the mouse head

To demonstrate the effectiveness of our ARCHmap and blockface-VISoR approach in studying the PNS, we first analyzed the previously inaccessible mesoscale structure of the mouse head using 3D reconstructions of the Thy1-EGFP mouse with DL649-labeled vasculature (Figure 2A-B). This dataset clearly revealed the fine architecture of cranial nerves, including individual fibers such as those in the trochlear nerve (CN IV), facial nerve (CN VII), and trigeminal nerve (CN V) (Figure 2C-2D).

**Figure 2.**
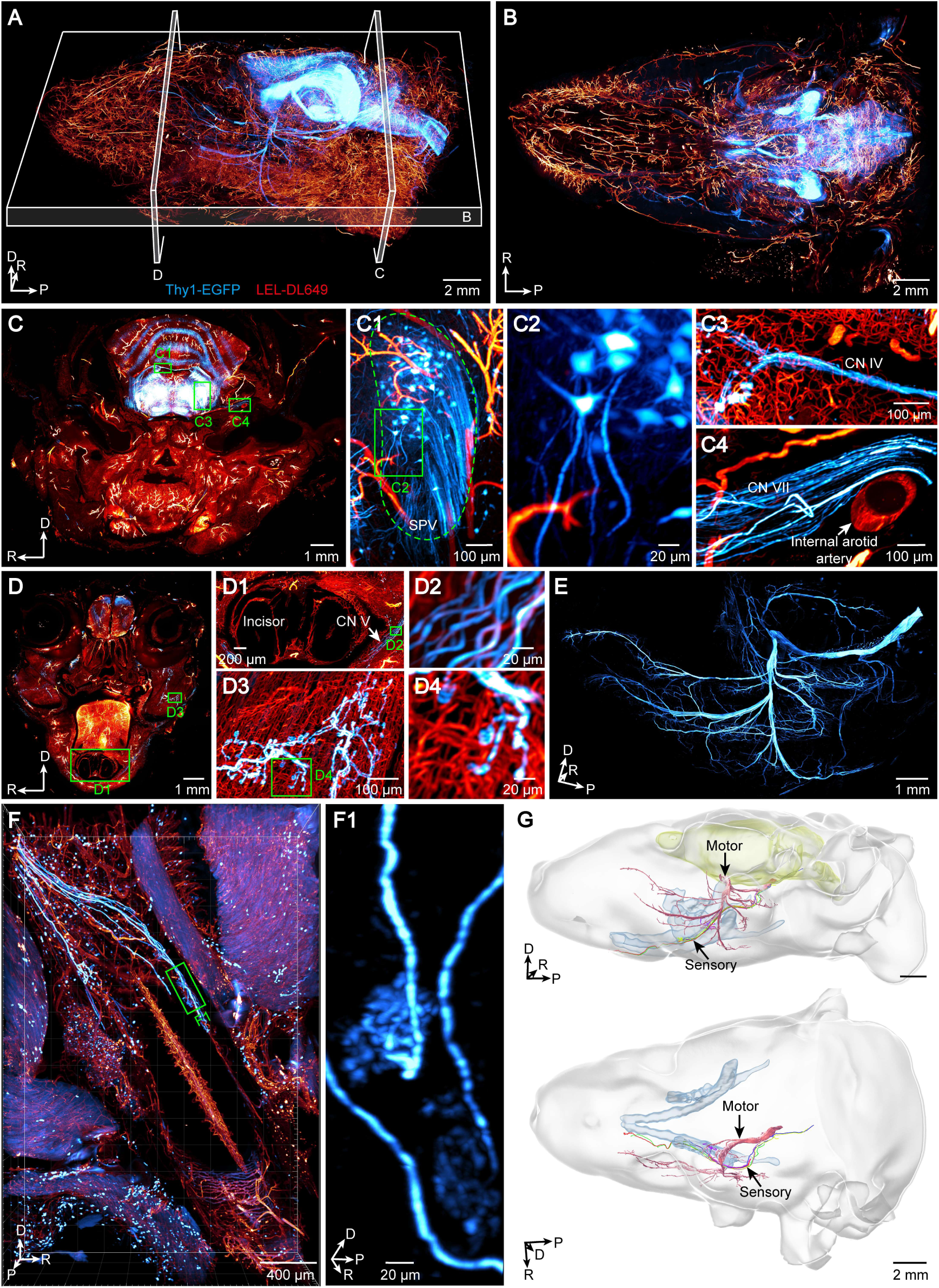
Visualization of cranial nerves. (A) 3D view of nerve and vasculature in a Thy1-EGFP mouse head. Red hot, LEL-DL649; cyan hot, Thy1-EGFP. (B) Maximum intensity projection of a 1-mm horizontal block in (A). (C, D) Maximum intensity projection of 100-μm coronal sections in (A). (C1-C4) Magnification of the spinal trigeminal nucleus (SPV) (C1, zoomed-in view in C2), the trochlear nerve (CN IV) (C3), and the facial nerve (CN VII) (C4) in (C). (D1-D4) Magnified sensory nerve fibers of the periodontium (D1, zoomed-in view in D2), and trigeminal motor terminals of the masseter muscle (D3, zoomed-in view in D4) in (D). (E) 3D view of the motor portion of the CN V segmented from (A). (F, F1) Sensory afferents of the CN V to the periodontium. (F1) Magnified view of the sensory endings in (F) from another perspective. (G) Single-fiber tracing of the trigeminal sensory nerves innervating the periodontium. Light blue surface, mandible; light yellow surface, brain; pink surface, motor portion of the CN V. Arrows indicate the diverse routes of sensory and motor components of the CN V. Orientation of images: R: right; P: posterior; D: dorsal.

Taking advantage of the uniformly high resolution of the dataset, we conducted a compressive and detailed analysis of the EGFP-expressing CN V. The motor portion of CN V displayed numerous thick fascicles and dense arborizations, which innervated the cephalon-facial muscles via widespread claw-like neuromuscular junctions (Figure 2D-2E). The sensory portion of CN V also exhibited thick, though sparsely labeled, sensory fibers in the Thy1-EGFP mice. These sensory nerves, originating from the semilunar ganglion, traversed the cavernous sinus alongside other cranial nerves (Figure S4A-S4B). Notably, we observed one sensory axonal fascicle of CN V innervating the periodontium of the incisor and molar teeth (Figure 2F-2G). Tracing these individual sensory nerves revealed a distinct pathway, separate from that of the motor components of CN V (Figure 2G).

### Reconstruction of single-axonal innervations from thoracic spinal nerves

The Thy1-EGFP mouse exhibits sparse expression of EGFP in spinal neurons, allowing us to distinguish individual fibers in the whole-body images acquired using the ARCHmap-blockface-VISoR workflow (Figure 3A-3C). We traced the long projections (up to approximately 8 cm) of thoracic motor neurons in the spinal cord and sensory neurons in the dorsal root ganglia (DRG). This analysis revealed clear visibility of distal structures, including neuromuscular junctions and sensory nerve endings such as free endings, lanceolate endings, and muscle spindles (Figure 3D-3I, S4C-S4G, and Video S3B-S3C). Being able to identify the complete structure of individual motor and sensory neurons, we observed that in the ventral rami, all motor fibers and some sensory fibers projected along the intercostal spaces toward the chest and the abdominal wall (Figure 3D and S4C-S4E). Notably, more than half of the sensory nerves in the ventral rami exhibited large V-shaped turns (Figure 3E). In the dorsal rami of the thoracic spinal nerves, the morphological differences between sensory and motor nerves became more pronounced: all sensory nerves displayed long axonal fibers with multiple zigzag-like turns (Figure 3F-3G and S4F), while the thoracic motor fibers were much shorter and exhibited dense arborizations (Figure 3H-3I).

**Figure 3.**
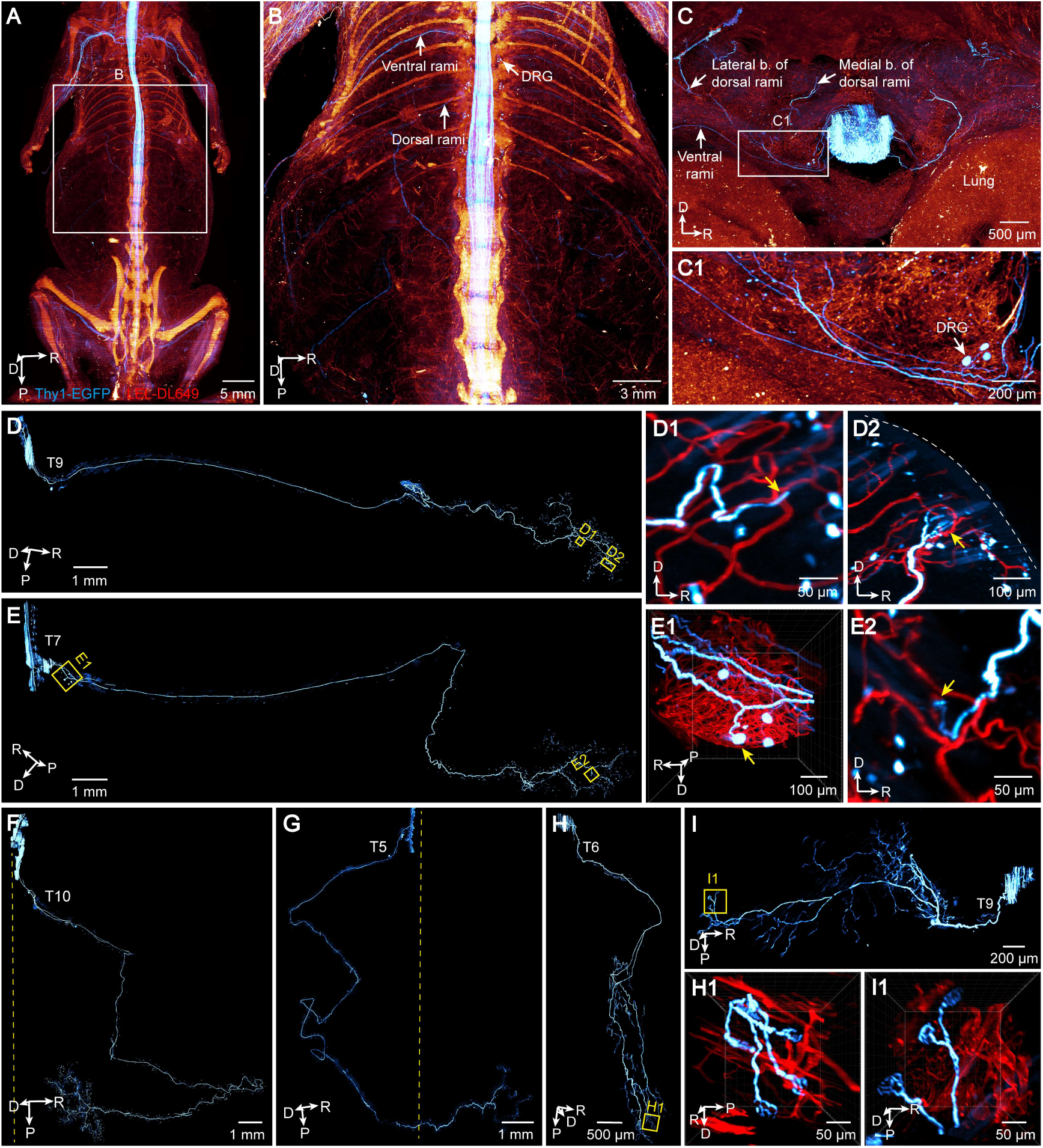
Visualization of individual thoracic spinal sensory and motor neurons. (A) 3D view of spinal nerves and vasculature in the body of a Thy1-EGFP mouse. Red hot, LEL-DL649; cyan hot, Thy1-EGFP. (B) Magnified 3D view of thoracic spinal nerves in (A). (C, C1) Maximum intensity projection of 1 mm-thick stacks showing rami and branches of the spinal nerves at the T4 segment (zoomed-in view in C1). (D-E) Representative anatomical morphology of individual spinal sensory neurons in the ventral rami characterized by forming no sharp turns (D) and sharp turns (E). (E) is viewed in a horizontal mirror-reversed perspective. (D1-D2, E1-E2) Magnified regions in (D) and (E) showing sensory terminals, including free nerve endings (yellow arrows in (D1) and (E2)) and the lanceolate endings (yellow arrow in D2), as well as the pseudounipolar neuron (yellow arrow in E1) in the dorsal root ganglion. White dashed lines in (D2) indicate skin surface. (E1) displays another perspective. (F-G) Different anatomical morphologies of individual spinal sensory neurons in the dorsal rami innervating the ipsilateral (F) and contralateral (G) skin. Yellow dashed lines indicate the midspinal line. (H-I) Different anatomical morphologies of individual spinal motor neurons in the medial (H) and lateral branch (I) of the dorsal rami. (H1, I1) Magnified neuromuscular junction regions in (H) and (I). (H1) displays another perspective. Orientation of images: R: right; P: posterior; D: dorsal.

From two reconstructed Thy1-EGFP mice, we traced the peripheral projections of 191 EGFP-expressing spinal neurons from segments T2 to T13. This included 66 motor and 66 sensory neurons from the ventral rami, as well as 30 motor and 29 sensory neurons from the dorsal rami (Figure 4A-4B and Video S3A). The compiled projection maps revealed distinct innervation patterns of different spinal neuron types in relation to their territorial segments (Figure 4A-4D). Quantitative analysis further demonstrated variability in the length and branching extent of individual spinal motor and sensory neurons in both the ventral and dorsal rami (Figure 4E-4J). For instance, motor fibers in the ventral rami exhibited a clear trend of segment-dependent increases in main fiber length from T2 to T13, whereas sensory fibers displayed relatively uniform lengths across segments (Figure 4F). Notably, motor fibers in the dorsal rami exhibited the most extensive branching, with a segment-dependent increase in both branching number and total branch length from T2 to T10 (Figure 4G-4J).

**Figure 4.**
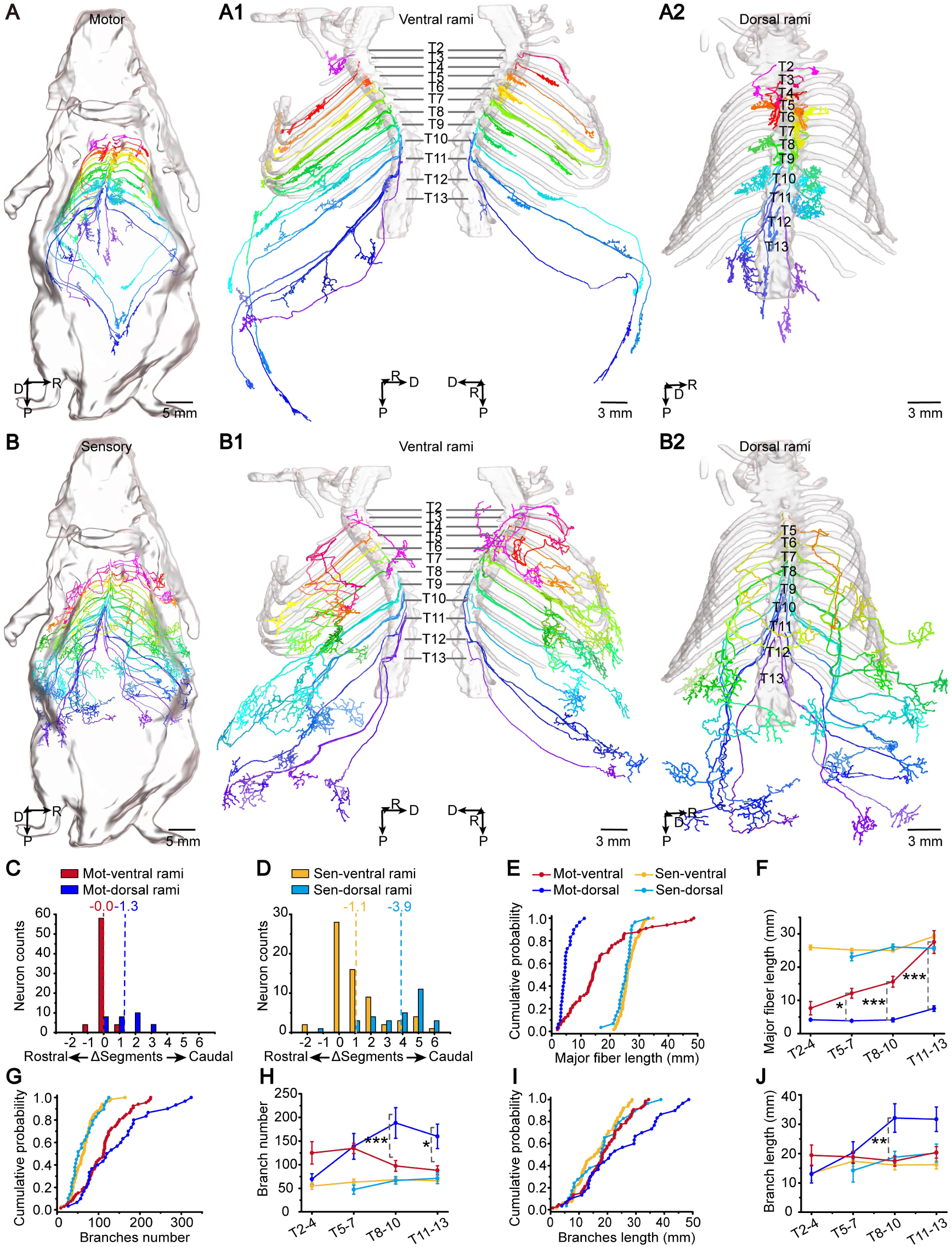
Anatomical comparison and analysis of individual thoracic spinal sensory and motor neurons. (A-B) Whole-body tracing of individual thoracic spinal motor (A) and sensory (B) neurons from two Thy1-EGFP mice aligned by thoracic skeleton. (A1-A2) 66 and 30 thoracic spinal motor neurons traced along the ventral (A1) and dorsal rami (A2), respectively. White surface, spine and ribs. 41 and 25 neurons from the left and right ventral rami, respectively; 14 and 16 neurons from the left and right dorsal rami, respectively. (B1-B2) 68 and 29 thoracic spinal sensory neurons in the ventral (B1) and dorsal rami (B2) of the spinal nerves, respectively. 39 (including 2 muscles spindle neurons that were excluded in further analysis) and 29 neurons from the left and right ventral rami, respectively; 12 and 17 neurons from the left and right dorsal rami, respectively. (C-J) Quantitative analysis and comparison of the morphological characteristics of spinal sensory and motor neurons. (C-D) Quantification of crossed spinal segments for motor (C) and (D) sensory nerves in both rami. Numbers above the dashed lines indicate averaged crossed segments. (E, G, I) Cumulative distribution of major fiber length (E), branches number (G), and branch length (I). 66 sensory neurons were of the ventral rami, 29 sensory neurons were of the dorsal rami, 66 motor neurons were of the ventral rami, and 30 motor neurons were of the dorsal rami from 2 mice. (F, H, J) Quantification of major fiber length (F), branch number (H), and branch length (J) at different thoracic segmental ranges. 6 to 23 neurons for each group; three-way ANOVA followed by Bonferroni post hoc test. (F) *P = 0.019 at T5-7, ***P = 3.37 × 10-4 at T8-10, ***P = 8.55 × 10-13 at T11-13. (H) *P = 0.026, ***P = 4.98 × 10-4. (J) **P = 0.0080. Data are presented as mean ± SEM. Orientation of images: R: right; P: posterior; D: dorsal.

The innervation patterns of individual spinal neurons, revealed through our high-resolution imaging of the whole mouse body, demonstrate cell type- and location-dependent diversity that traditional anatomical descriptions of the intercostal or subcostal nerves do not capture.^48^ Efficient mapping of peripheral neuronal projections in mice, utilizing sparse labeling in combination with the ARCHmap and blockface-VISoR systems, may facilitate the construction of a mesoscopic whole-body connectome and reveal crucial functional insights, paralleling the development of the still-growing mesoscopic whole-brain connectome.^13,49,50^

### Visualization of immunolabeled sympathetic system

To evaluate the compatibility of our approach with immunolabeling, we integrated an antibody against tyrosine hydroxylase (TH) into our ARCHmap workflow (see Methods) to image the sympathetic nervous system. This system, widely distributed throughout the body, regulates the physiological activities of organs and vasculature.^51–56^ Although previous studies have examined the fine architecture of sympathetic nerves in isolated rodent organs or partial primate organs,^57–61^ a comprehensive high-resolution map of sympathetic projections to all organs in the body remains absent. Using blockface-VISoR imaging, we observed extensively labeled sympathetic nerves throughout the entire mouse body (Figure 5A and Video S4A). We clearly visualized the sympathetic chain and plexuses, as well as fine nerve branches connecting to adjacent paravertebral ganglia and prevertebral ganglia (Figure 5B-5D, S5A-S5B, and Video S4B). The efficiency and specificity of our whole-body immunolabeling strategy were further confirmed by strong anti-TH signals in presumed TH^+^ catecholamine neurons and fibers within the brain and spinal cord (Figure 5E), and by their colocalization with genetically labeled peripheral sympathetic postganglia and fibers (Figure 5C-D and S5B-S5C).

**Figure 5.**
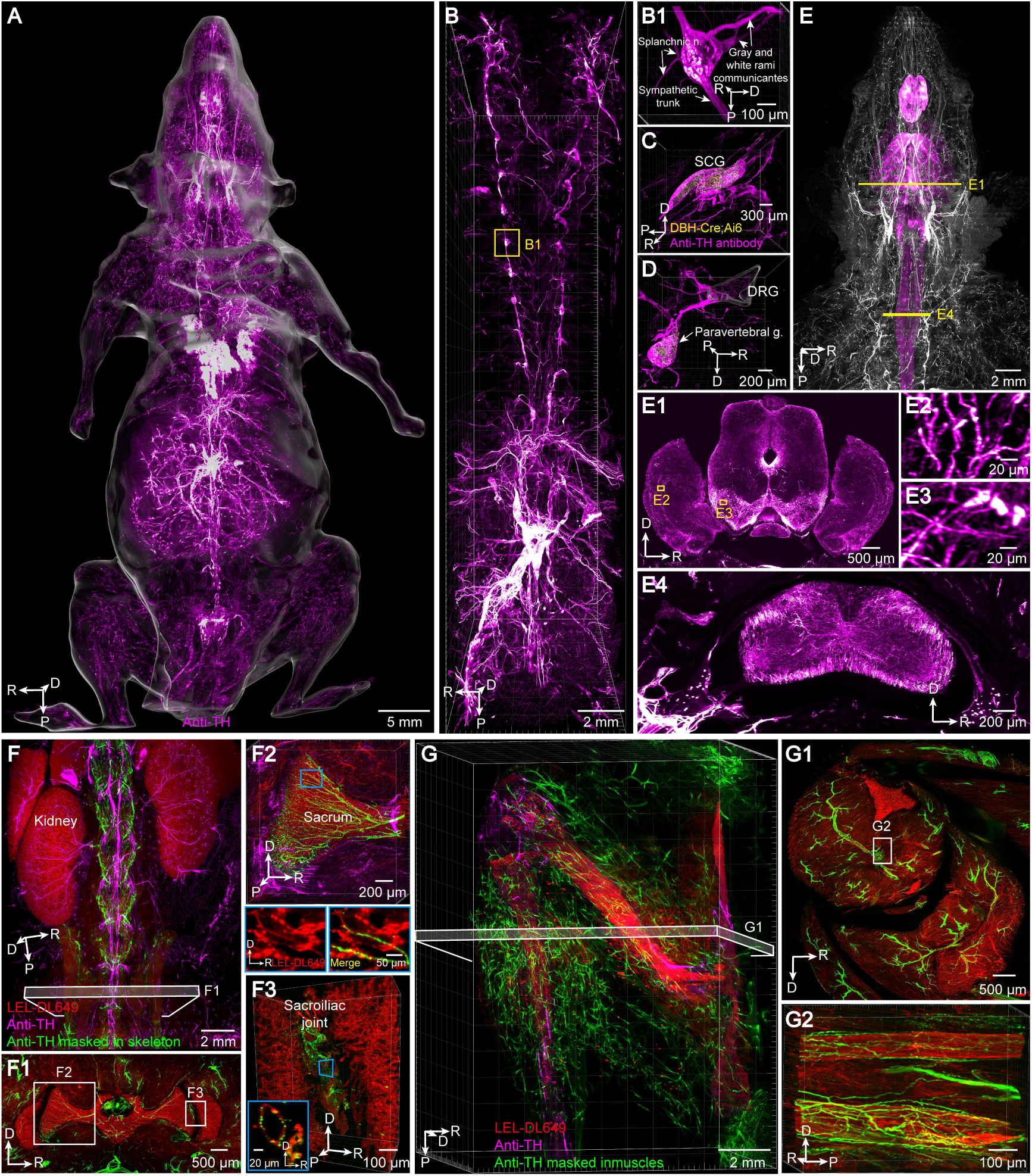
Visualization of immunostained sympathetic nerves in whole adult mouse. (A) 3D view of the sympathetic nervous system in an adult mouse via whole-body immunostaining with the anti-TH antibody. (B) Ventral view of the segmented sympathetic chains and plexuses of the mouse. (B1) Magnified PVG and connected branches in (B). (C-D) 3D views of immunostained SCG (C) and thoracic paravertebral ganglia (D) in a DBH-Cre;Ai6 mouse with endogenous ZsGreen expressions. (E) 3D view of sympathetic nerves in a mouse head with the CNS segmented in magenta pseudo-color. (E1-E4) Representative images in (E) showing uniform TH staining of the brain (E1) and the spinal cord (E4). Images are the maximum intensity projection of 100- and 400-μm-thick stacks for (E1) and (E4), respectively. (E2-E3) Enlargement of TH+ somas and fibers in (E1). (F) 3D view of sympathetic nerves and vasculature in the spine. (F1) Maximum intensity projection of 400-μm coronal sections in (F) showing sympathetic nerves in the sacral vertebra. (F2-F3) Magnified 3D view of sympathetic nerves in the sacral vertebra (F2) and the sacroiliac joint space (F3). Magnified inserts are the maximum intensity projection of 10-μm stacks showing perivascular sympathetic nerves in the skeleton. (G) 3D view of sympathetic nerves and vasculature in the left leg. (G1-G2) Maximum intensity projection of 400-μm stacks in (G) showing perivascular sympathetic nerves in skeletal muscles of the leg. (G2) 3D magnification of circumjacent regions in (G1) from another perspective. Orientation of images: R: right; P: posterior; D: dorsal.

By combining uniformly high-resolution visualization of the sympathetic system with lectin labeling, we further evaluated how sympathetic nerves interact with blood vessels throughout the mouse body. For example, when zooming into the lumbosacral skeleton, we found that sympathetic nerves traveled alongside blood vessels in the sacrum and sacroiliac joint (Figure 5F). Additionally, we observed dense sympathetic innervations intertwining with surrounding vasculature within the leg muscles (Figure 5G). This perivascular pattern of sympathetic projections was consistently observed in various regions of the body trunk and limbs.

Whole-body visualization of sympathetic nerves and their fine branches allows for systematic mapping of their architecture, revealing distinct projection routes and innervation patterns to various internal organs. In the cranial regions, a sympathetic nerve from the superior cervical ganglion (SCG) bifurcates to innervate different salivary glands (Figure S5D); another projected to the cochlea, forming a pinwheel-like arborization structure (Figure 6A and Video S5A). In the thoracic segment, sympathetic nerves from multiple paravertebral chain ganglia project to the heart where they extend branches (Figure S6A). Within the abdomen, sympathetic nerves projecting to the kidneys from the prevertebral ganglia (PVG) penetrate the renal parenchyma, predominantly targeting the renal cortex and twining around blood vessels (Figure 6B-6C, S6B-S6C and Video S5B), with minuscule terminals enveloping glomeruli clearly visible (Figure 6B1). Additionally, sympathetic innervations to the spleen, liver, and pancreas exhibit clear perivascular patterns (Figure 6D, S6D-S6E, and Video S5C-S5D). In contrast, sympathetic fibers in the stomach and intestine form meshwork-like structures along the gastrointestinal wall, largely nonoverlapping with the vasculature (Figure S6F-S6G). This pattern may relate to their role in regulating gastrointestinal smooth muscle activity.^62^

**Figure 6.**
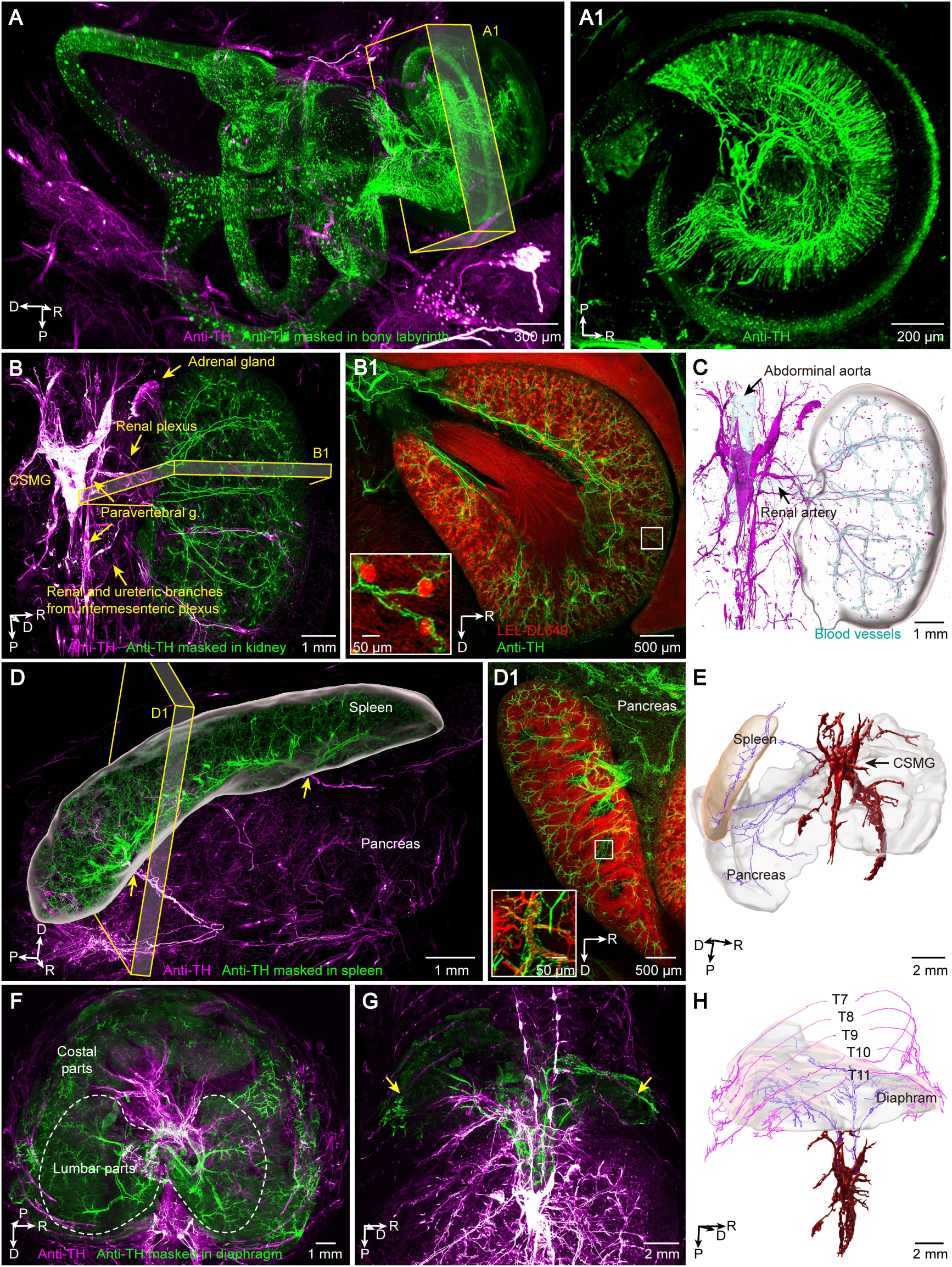
Visualization of the sympathetic innervations to organs. (A) 3D view of sympathetic nerves in the left bony labyrinth segmented in green pseudo-color. (A1) Maximum intensity projection of a 500-μm x-stack in (A) showing radial-like sympathetic networks in the cochlea. (B) 3D view of the sympathetic innervations to the kidney segmented in green. (B1) Maximum intensity projection of 400-μm coronal sections in (B) showing dense sympathetic networks and vasculature in the renal cortex. Magnified insert is the maximum intensity projection of 100-μm coronal sections showing sympathetic innervations to the glomeruli. (C) Normal shading view of major sympathetic branches surrounding blood vessels (light cyan surface) in the kidney (white surface). (D) 3D view of the sympathetic innervations to the spleen segmented in green. (D1) Maximum intensity projection of 400-μm coronal sections in (D) showing sympathetic networks and vasculature in the spleen. Magnified insert is the maximum intensity projection of 100-μm coronal sections showing sympathetic innervations to the white pulp of spleen. (E) Axonal tracing showing shared sympathetic innervations to the spleen and pancreas from the CSMG. Orange surface, spleen; white surface, pancreas; red surface, CSMG. (F-G) 3D view of sympathetic innervations to the diaphragm segmented in green. White dashed lines indicate the border of the diaphragm lumbar parts. Arrows in (G) indicate the entering the points of sympathetic nerves from thoracic paravertebral chain ganglia to the costal diaphragm. (H) Fiber tract tracing of sympathetic innervations to the diaphragm from both the CSMG (purple lines) and the thoracic paravertebral ganglia (magenta lines). White surface, diaphragm. Orientation of images: R: right; P: posterior; D: dorsal.

Tracing the projection routes from the coeliac-superior mesenteric ganglion (CSMG) revealed distinct pathways to visceral organs, including the spleen, pancreas, liver, and stomach (Figure 6E and S6H-S6I). These routes largely align with those observed in humans. We then focused on the diaphragm, a somatic muscle essential for breathing. In addition to motor inputs,^63–65^ the diaphragm also receives sympathetic innervations.^66,67^ However, the sources of these sympathetic innervations and their regional specificity were unclear. With high-resolution mapping, we identified distinct innervation zones in the diaphragm’s muscular portion, each receiving sympathetic inputs from different sources (Figure 6F-6G). Sympathetic clusters in the lumbar diaphragm traced back to both the CSMG and phrenic ganglion via the crura, while clusters in the costal diaphragm connected to thoracic paravertebral chain ganglia (Figure 6H and S6J1-S6J2). This mesoscopic mapping provides an unambiguous and precise description of sympathetic projections, surpassing the capabilities of conventional anatomical methods.

### Visualization of virus labeled vagus nerves and single fibers

To assess our approach’s capability for mapping peripheral nerves using viral labeling, which is commonly applied in brain connectomics, we injected AAV-EF1α-tdTomato into the left and right dorsal motor nuclei of the vagus nerve (DMV) and AAV-CAG-EYFP into the left vagal ganglia, including the nodose ganglia (NG) and jugular ganglia (JG) (Figure S7A-S7B). Whole-body imaging enabled us to trace the motor and sensory components of the vagus nerve and their innervation routes to various organs (Figure 7A and S7C-S7D). At the cervical level, sensory and motor fibers of the vagal trunk remained spatially separated, but they became increasingly intermingled in the distal segments (Figure 7B and 7C1-7C4). At the thoracic level, they also received the contralateral nerve fascicles, showing a more complexed distribution pattern within the branch (Figure 7C4). This provides a high-resolution view of the anatomical organization and mixing of sensory and motor fascicles, complementing previous micro-computed tomography imaging studies.^68^

**Figure 7.**
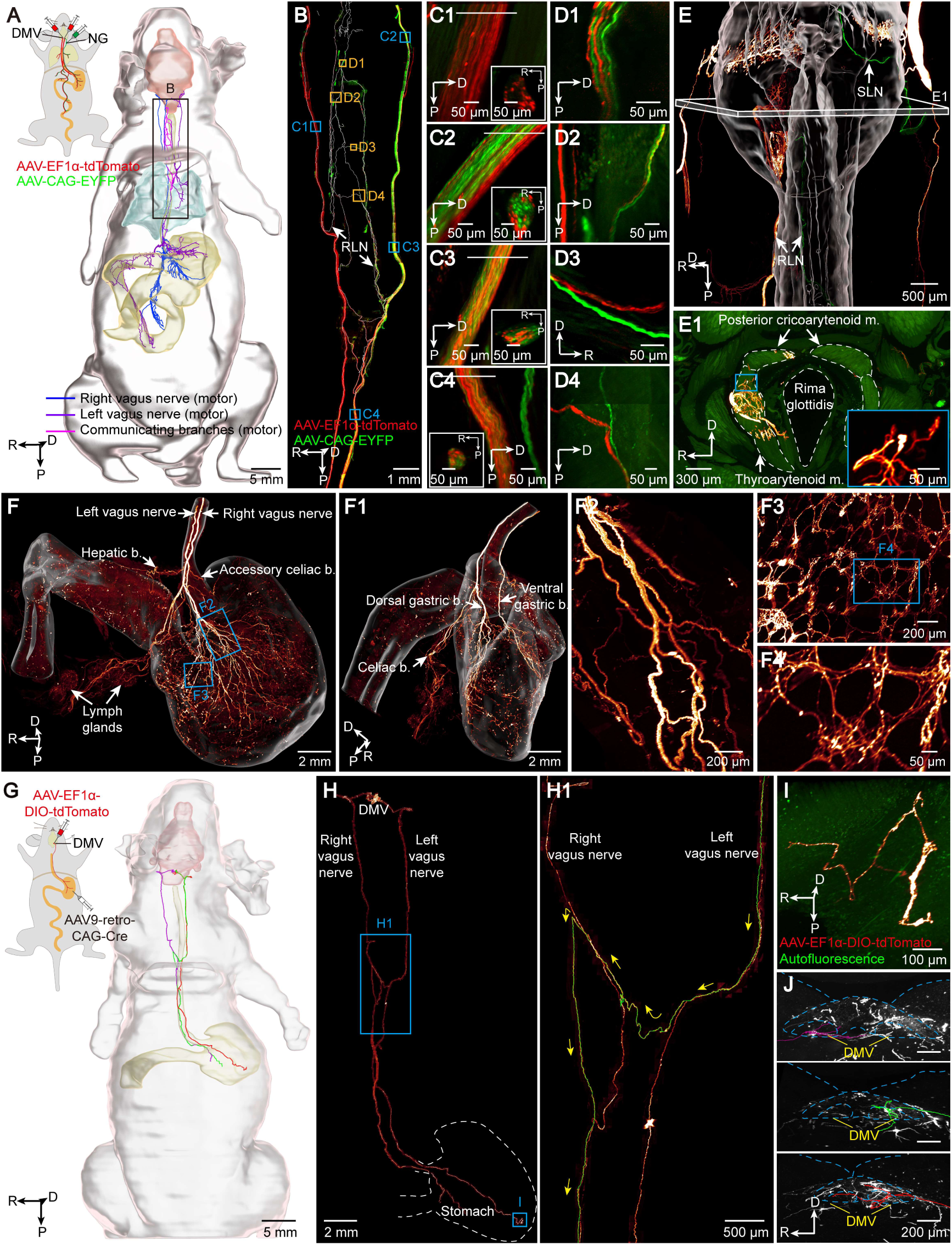
Visualization and individual neuron tracing of the viral-labeled vagus nerve. (A) Whole-body tracing of the dense-labeled visceral motor fibers of the vagus nerve. Blue lines, left vagus nerve; purple lines, right vagus nerve; magenta lines, communicating branches between the left and right vagus nerves. Viral labeling procedures are shown in the schematic. Pink surface, brain; cyan surface, lung; yellow surface, digestive tract. (B) 3D view of the segmented supradiaphragmatic vagus nerves, including both general visceral sensory and motor fibers. (C1-C4) Magnified vagal trunk in (B) showing spatial distribution of sensory and motor fibers in the vagal trunks. Insert panels are the maximum intensity projections of 5-μm z-stacks indicated in the cross section of (C1-C4). (D1-D4) Magnified view of communicating branches between the left and right RLN in the tracheoesophageal and pulmonary plexus in (B). (D3) displays another perspective. (E) 3D view of the vagal sensory and motor innervations to the larynx. White surface indicates the trachea, esophagus, and surrounding laryngeal muscles. (E1) Representative images showing vagal innervation to the laryngeal muscles. Images are maximum intensity projection of 100-μm z-stacks in (E). Magnified insert indicates vagal motor endings. (F, F1) 3D view of vagal innervations to the stomach and duodenum. (F1) Side view of (F) showing vagal branches. (F2-F4) Magnified superficial view of the gastric branches (F2) and myenteric plexus (F3, zoomed-in view in (F4)) in (F). (G) Individual neuron tracing of three stomach-projecting vagus nerves across the whole body. Viral labeling procedure of the injection of AAV9-retro-CAG-Cre into the stomach and AAV-CAG-DIO-EGFP into the DMV are shown in the schematic. (H, H1) 3D reconstruction of entire projecting routes of three individual vagus neurons. (H1) Magnified supradiaphragmatic segment of the vagus neurons. (I) Magnified axonal terminals of a vagal neuron in the gastric wall in (H). (J) Maximum intensity projection of 64-μm z-stacks showing tracing of three vagal neurons back to the DMV. Orientation of images: R: right; P: posterior; D: dorsal.

Based on this mapping, we observed multiple interconnections between the left and right recurrent laryngeal nerves (RLNs) and between the left and right vagal trunks (Figure 7B and 7D1-7D4). By using EYFP labeling to target left vagal sensory neurons, we identified fibers from the left RLN crossing at various points to the right side to innervate the larynx in two of the three mice examined (Figure 7D-7E and S7E). Additionally, tdTomato-labeled motor fibers in the RLNs displayed extensive crossover, forming dense arborizations within the laryngeal muscles (Figure 7D-7E and S7E). The features of these previously unrecognized crossover fibers resemble those seen in the supradiaphragmatic communicating branches of the esophageal plexus, which connect the left and right vagal trunks (Figure S7C-S7D). However, there are no crossing-over fibers above the vocal cord level (Figure S7E).

In all three virus-labeled mice, we observed that the labeled vagal motor and sensory fibers exclusively innervated the pharynx, larynx, esophagus, lungs, stomach, and intestines (Figure 7A, 7E-7F, and S7F-S7H). In the abdomen, 3D reconstruction revealed dense innervation of the gastric branches from the left and right motor vagal trunks, targeting the ventral and dorsal walls of the stomach, respectively (Figure 7F, 7F1, and Video S6B). Zoom-in views showed a meshwork of fasciculated axons forming the myenteric plexus (Figure 7F2-7F4).^69^

To further assess the capability of our approach for single-fiber tracing, we sparsely labeled stomach-projecting visceral motor fibers of the vagus nerve by injecting AAV9-retro-CAG-Cre into the stomach^70^ and AAV2/9-EF1α-DIO-tdTomato into the DMV (Figure 7G). This approach resulted in strong labeling of three individual vagus neurons in the DMV and their fibers (Figure 7H), allowing for clear tracing. Two of these fibers traveled along ipsilateral vagal trunks to innervate the ventral and dorsal sides of the stomach. The third fiber, originating from the left DMV, crossed to the contralateral side at a supradiaphragmatic location. It then traveled near the right vagal fiber but surprisingly in the rostral direction, then made a sharp turn, and returned caudally to terminate in the dorsal stomach (Figure 7H1 and 7I-7J).

We traced six individual vagal sensory neurons from another mouse that received AAV9-retro-CAG-Cre injections into the tail vein and AAV2/9-CAG-FLEX-EGFP injections into the left NG (Figure S7I). One neuron crossed to the contralateral side via the communicating branch and projected to the dorsal stomach, while the other five innervated the left bronchi. Four of these fibers took V-shaped turns just before entering the lungs (Figure S7I-S7K). Notably, the individual motor and sensory vagal neurons traced in this study were all unbranched in the vagal trunk, indicating a straightforward one-to-one projection pattern to the target organs. However, the projection routes of these neurons exhibited complexity, featuring frequent contralateral crossings and intricate turns both mid-route and near their targets.

## Discussion

In this study, we combined series thick sectioning with the blockface-VISoR technique to image the cleared whole mouse body. By acquiring oblique image series illuminated by a synchronized scanning light beam while continuously moving the sample, the VISoR technique achieves ultrahigh speed without motion blur. Additionally, the blockface approach ensures uniformly high resolution throughout the entire body by imaging hundreds of microns into the cut surface of the cleared sample. Using our prototype system, imaging an adult mouse body at a voxel size of 1 × 1 × 2.5 µm^3^ (∼70 TB data) takes approximately 40 hours for each fluorescence channel. We can achieve higher throughput (in terms of data rate) by employing two or more cameras to simultaneously acquire multi-channel images or by using faster cameras. Moreover, our system can be readily scaled to image larger animal samples.

Achieving high-quality blockface-VISoR imaging depends critically on the preparation of a cleared whole mouse body. Using ARCHmap, we optimized several key factors, including tissue transparency, preservation of antibody labeling and fluorescent protein signals, minimization of tissue volume changes, and the mechanical properties of embedding. This balanced approach enables fluorescence imaging of the entire mouse body at uniform subcellular resolution that is difficult to achieve using existing methods. The blockface-VISoR system can also be applied to solvent-cleared samples, such as those prepared using TESOS or DISCO depending on the requirements of specific applications.^23–25,33^ As long as the cut surface remains smooth, these samples can be imaged effectively. Additionally, the high mechanical strength of solvent-cleared samples minimizes deformation during sectioning, simplifying the reconstruction of blockface-VISoR images.

Our technology and workflow effectively demonstrate the ability to visualize the PNS at both large and fine scales. The high-quality imaging and reconstruction enable us to observe previously inaccessible structures and uncover new features of the PNS. One particularly notable finding is the body-wide architecture of the sympathetic nervous system. Classical anatomy has long recognized that sympathetic nerves form mesh-like configurations around arteries.^51^ However, the prevalence of this perivascular pattern throughout the entire body at subcellular resolution, or mesoscale, has not been systematically examined due to technical limitations. Our observations reveal the widespread presence of perivascular meshwork sympathetic nerve fibers in the head and neck, torso, limbs and most internal organs, with the exception of the stomach and intestine, where many sympathetic fibers do not follow blood vessels. This mesoscale mapping of sympathetic architecture holds clinical relevance for procedures such as superficial neural stimulation and radiofrequency ablation, used to treat conditions like pain and hypertension.^71–73^

Another crucial aspect of our method is its ability to identify and trace individual nerve fibers throughout the entire body. This capability allows us to qualitatively visualize the complete projection routes of individual spinal sensory and motor neurons and to reveal distinct cell type- and rami-specific arborization features. Additionally, it enables an unambiguous evaluation of how vagus nerves innervate various visceral organs, particularly whether a single vagal neuron can send fiber branches to multiple organs.^74–76^ Using a sparse viral labeling approach, we traced the complete fibers of nine individual vagal neurons targeting the stomach or lungs. All these fibers were unbranched before entering their respective targets, indicating a prevalent one-to-one organ-innervating pattern for the general visceral fibers of the vagus nerve. It is reasonable to further presume that the DMV may be of multiple sub-zonings for various organs.

The ARCHmap-blockface-VISoR technique presented in this study offers a high-efficiency, high-resolution imaging method for the whole mouse that is compatible with diverse labeling strategies. This method extends mesoscale connectomic mapping from the brain to the entire nervous system. Beyond the nervous system, our approach facilitates system-level analysis of interactions between various cell types and tissues, provided they are properly labeled. We anticipate that such analyses will not only address long-standing questions in neuroscience but also provide valuable insights into broader areas such as developmental biology, comparative anatomy, and biomedical research in general.

### Limitations of the study

The lateral optical resolution of our system depends on the higher-numerical aperture (NA) of the imaging objective. However, the oblique imaging configuration of VISoR restricts the selection of objectives due to geometric constraints, precluding those with the highest NA. The axial resolution varies based on the thickness of the illuminating light beam, which is typically several micrometers. To achieve higher resolution, one can collect tissue sections for re-imaging using other methods, such as confocal and super-resolution microscopy,^77^ but only for specific regions of interest. Alternatively, one may also take computational approaches such as deep learning neural networks trained with high-resolution data, to achieve higher and isotropic resolution.^78,79^

The blockface-VISoR with ARCHmap workflow enables uniform fluorescence imaging throughout the mouse body. However, in some deep adipose tissues, we still see occasional blurry regions caused by inadequate delipidation. To mitigate this problem, further improvement in the tissue clearing protocol is needed. One possibility is to integrate solvent-based delipidation in the workflow when sample shrinkage does not compromise the structures of interest.^80,81^ Additionally, although tissue sections generated during whole-body imaging can be collected for reimaging, they were hard to be further re-stained using new antibodies, possibly due to blocking of antigenic sites during hydrogel embedding. Ideally, future optimization of embedding methods should aim for compatibility with re-staining to further expand the capability the blockface-VISoR system.

## Lead contact

Further information and requests for resources should be directed to and will be fulfilled by the Lead Contact, Guo-Qiang Bi.

## Acknowledgements

We thank Yingrui Li for helping with video making of imaging setup. This study was supported by the National Key Research and Development Program of China (grant No. 2019YFA0801601 to G.B. and P.L.), the National Natural Science Foundation of China (grant No. 32071039 to Q.Z. and No. 92253301 to G.B.), National Natural Science Foundation of China-Guangdong Joint Fund (No. U20A6005 to P.L.), the STI2030-Major Projects (grant No. 2021ZD0204400 to C.X.), and the Strategic Priority Research Program of Chinese Academy of Science (grant No. XDB32030200 to G.B.), the Fundamental Research Funds for the Central Universities (grant No. WK9990000138 to M.S.), and the Natural Science Foundation of Anhui Province (grant No. 2408085QC086 to M.S.).

## Author contributions

G.B., C.X., P.L., Q.Z., and M.S. conceptualized the project. M.S. and C.X. designed and interpreted the experiments under the supervision of G.B., P.L., and Q.Z.. C.X. and Q.Z. designed and constructed the imaging instrument. C.X. and Z.Z. designed and constructed the vibratome. M.S., Y.Y., and C.X. established the whole-body clearing and embedding pipeline. M.S., Y.Y., and M.W. performed whole-body immunostaining or viral labeling. Y.Y., M.S., Y.G., and Z.S. performed organ clearing tests. L.D., C.X., and C.Y. developed the software for image acquisition. Y.Y., M.W., M.S. and C.X. performed whole-body imaging and acquired data. Q.Y., L.D., R.L., and C.Y. wrote the codes for whole-body reconstruction or image processing under the supervision of C.X. and Q.Z.. Q.Y. and C.X. reconstructed the whole-body images. M.S., Y.Y., M.W., F.D., and X.Z. performed individual neuron tracing and analyzed data. L.Q., Y.L., and H.H. performed 3D alignment. F.X., Z.X.Z., and P.W. provided the AAV9-retro virus. K.Z. designed and generated 3D printing molds for embedding. M.S., C.X., Y.Y., Z.Z and C.X. designed and generated the figures and videos. H.X., L.D., and Q.Y. built the data browsing webpages. M.Z., Y.O.C., and Z.R. provided neuroanatomical insights. M.S., C.X., Q.Z., P.L. and G.B. wrote the manuscript with feedback from M.Z. and all other authors.

## Data and code availability

Image data and all original image stacks will be accessible to all scientists as of the date of publication from the lead contact upon request. All original code is deposited at GitHub and will be publicly available as of the date of publication. Any additional information required to reanalyze the data reported in this paper is available from the lead contact upon request.

## Competing interests

The authors declare no competing interests.

## Additional information

2D maximum intensity projection of consecutive mouse sections can be viewed at a dedicated web page: https://mesoanatomy.org/mesomouse/meso-example. The website provides image browsing at different zoom-in levels.

## Methods

### Mice

All experimental procedures in this study followed international guidelines and protocols approved by the Institutional Animal Care and Use Committee of the University of Science and Technology of China. We used adult male and female mice from the following strains: C57BL/6J (stock#: 000664), Thy1-EGFP (stock#: 007788), CaMKIIα-Cre (stock#: 005359), vGAT-Cre (stock#: 028862), DBH-Cre (stock#: 036734-UCD), Ai6 (stock#: 007906), and Ai14 (stock#: 007914). The ages of the mice ranged from 8 to 20 weeks. We housed the mice in a temperature-controlled (22 °C) colony room with a 12-hour light/dark cycle and provided *ad-libitum* access to food and water, except during periods of food restriction as described below.

### Surgery and virus injection

To achieve dense viral labeling of the visceral sensory vagus nerve, we anesthetized C57BL/6J mice with pentobarbital sodium (100 mg/kg, i.p.) and positioned them in a stereotaxic apparatus. We carefully exposed the left vagal ganglia through dissection. We administered a microinjection of 500 nl of purified and concentrated AAV carrying vectors expressing CAG-promoted EYFP (AAV2/9-CAG-EYFP, virus titer: 1.1×10^13^ v.g/ml). For viral labeling of the visceral motor vagus nerve, we removed the skull above the dorsal motor nucleus of the vagus nerve (DMV) using a dental drill and unilaterally microinjected 200 nl of AAV2/9-EF1α-tdTomato (virus titer: 2.5×10^12^ v.g/ml) at coordinates from bregma AP/ML/DV: −7.50 ∼ −7.90/0/-5.35 mm, using a syringe micropump at a rate of 40 nl/min. For sparse labeling of the vagus nerve, we injected 2-5 μl of AAV9-retro-CAG-Cre (virus titer: 8.9×10^12^ v.g/ml) into the front wall of the glandular stomach or the tail vein. Additionally, we microinjected 200 nl of AAV2/9-EF1α-DIO-tdTomato (virus titer: 2.1×10^12^ v.g/ml) or 300 nl of AAV2/9-CAG-FLEX-EGFP (virus titer: 1.1×10^13^ v.g/ml) into the left and right DMV or the left vagal ganglia, respectively. We allowed the mice to express the viral vectors for 2 to 3 months before sacrifice.

### Perfusion and Postfixation

*Perfusion*: For whole-body imaging, we fasted the mice and deprived them of water, while allowing free access to a 7% w/v polyethylene glycol electrolyte solution in 5% sucrose for 24 hours to empty the gastrointestinal tract. For both whole-organ tests and whole-body imaging, we deeply anesthetized the mice and exposed the thoracic cavity. We inserted a catheter into the left ventricle and secured it with a hemostatic clip at the cardiac apex. We then perfused the mice with 40 ml of saline at 37 °C, followed by 20 ml of 4% paraformaldehyde (PFA) in 0.1 M phosphate buffered saline (PBS) at room temperature through the cardiac catheter.

#### Postfixation

For *in vitro* clearing tests, we collected the brains and postfixed them in 4% PFA at 4 °C overnight following transcardial perfusion. For whole-body clearing experiments, we placed the catheterized mice in a chamber filled with 250 ml of 4% PFA and circulated the PFA intracardially for 24 hours at room temperature using a peristaltic pump at a rate of 5 ml/min. After postfixation, we rinsed the mice intracardially with 0.1 M PBS for 1 hour in the chamber, preparing them for further clearing procedures.

To evaluate the influence of the postfixation procedure on the permeation efficiency of reagents throughout the body, we prepared mouse samples using five postfixation protocols: passive postfixation in 4% PFA for 1 day at 4 °C; passive postfixation in 4% PFA for 1 day at room temperature; and intracardiac circulation with 4% PFA for 1 day at room temperature at three different rates: 1, 5, or 10 ml/min. Subsequently, we circulated all samples intracardially with 300 ml of 0.05% methylene blue dissolved in 0.1 M PBS for an additional day at room temperature at a rate of 5 ml/min. After methylene blue permeation, we removed the abdominal wall to expose the viscera and extracted the brains and spinal cords for further comparison.

### Vasculature labeling

To label the vasculature throughout the body, we anesthetized the mice and injected 0.2 ml of DyLight 649-conjugated lectin via the tail vein. Two minutes after the injection, we sacrificed the mice for transcardial perfusion as previously described. To enhance vascular labeling of the great vessels in Figure 2, we circulated the cleared mouse with an additional 0.2 ml of lectin dissolved in 40 ml of 0.1 M PBS for 2 hours at a rate of 5 ml/min before embedding.

### Whole-brain clearing

After three 2-hour washes in 0.1 M PBS, we transferred the postfixed brains to 50 ml conical tubes containing 40 ml of various clearing solutions. The clearing solutions used were as follows: PACT (8% w/v sodium dodecyl sulfate in 0.1 M PBS, pH 7.0),^84^ mPACT (8% w/v SDS, 0.5% w/v α-thioglycerol in 0.1 M PBS, pH 8.0),^40^ SeeNet (10% w/v SDC in a 200 mM boric acid-based buffer, pH 8.5),^41^ SHANEL (10% w/v CHAPS and 25% w/v N-methyldiethanolamine in dH_2_O),^85^ ScaleA2 (4 M urea, 10% w/v glycerol, and 0.1% w/v Triton X-100 in dH_2_O, pH 7.7),^43^ SUT (25% w/v urea, 15% v/v Triton X-100, and 8% w/v SDS in 0.1 M PBS, pH 7.4),^44^ FOCM (20% w/v urea, 30% w/v D-sorbitol, and 5% w/v glycerol in DMSO),^45^ CUBIC (25% w/v urea, 25% w/v quadrol, and 15% w/v Triton X-100 in dH_2_O),^46^ CUBIC-L (10% w/v Triton X-100 and 10% w/v N-butyldiethanolamine in dH_2_O),^47^ and CUBIC-LH (10% w/v Triton X-100 and 15% w/v N-butyldiethanolamine in dH_2_O). We incubated the brains in the aforementioned clearing reagents at 37 °C for 6 days with gentle shaking, refreshing the solutions every 2 days. We then washed the cleared samples three times with 0.1 M PBS at room temperature, each wash lasting 2 hours. In the final clearing procedure, we subjected the samples to refractive index (RI) matching in 40 ml of the Puclear RI-matching solution (RI of 1.52),^29^ which contains 50% w/w iohexol, 23% w/w urea, 11% w/w 2,2’,2”-nitrilotriethanol, and 16% w/w dH_2_O, at room temperature for an additional 3 days.

### Extracted tissue processing

We immersed the extracted cleared brains in an equal mixture of 20% w/v bovine serum albumin (BSA) and a 4% hydrogel monomer solution (HMS) containing 4% w/v paraformaldehyde (PFA), 4% w/v acrylamide, 0.05% w/v N,N′-methylenebisacrylamide, and 0.0025% w/v VA-044 in 0.1 M PBS for permeation at 4 °C for 24 hours. Afterward, we vacuumed the brains on ice for 20 minutes and polymerized them at 37 °C for 4 hours.

For fluorescence preservation tests, we extracted brains from CaMKIIα-Cre;Ai14 mice, embedded them, and sectioned them into 300-μm-thick slices using a vibratome. We subjected all slices to delipidation overnight with 4% SDS, followed by rinsing three times with 0.1 M PBS. The slices were then divided into several groups and immersed in various solutions, including 0.005% NaN₃ in 0.1 M PBS, mPACT (pH 8.0), SeeNet (pH 7.5), CUBIC (pH 8.5), CUBIC-L (pH 8.5), and CUBIC-LH (pH 8.5) for 4 days, and in 10% and 20% EDTA in water (pH 6.5 or 8.5) for two days each. After rinsing in 0.1 M PBS, we mounted the slices on glass slides. Before light-sheet imaging, we matched the brain slices with the Puclear RI-matching solution.^29^

We determined tissue size using bright-field images of unembedded 300-µm-thick brain slices. We processed the brain slices with various clearing and RI matching reagents. Initially, we immersed the samples in different clearing agents, including Scale, FOCM (30% w/v urea, 20% w/v D-sorbitol, and 5% w/v DMSO),^45^ PACT, mPACT, SeeNet, SUT, CUBIC, CUBIC-L, and CUBIC-LH, for one day. Most slices, except those treated with Scale and FOCM clearing reagents, were subsequently washed in 0.1 M PBS and treated with the corresponding RI-matching reagents for an additional day at 37 °C with gentle shaking. The RI-matching solutions included: 0.88 g/ml iohexol in 30 ml of base buffer (0.01% sodium azide and 0.1% Tween-20 in 0.02 M PB, pH 7.5) for PACT,^39^ mPACT,^40^ and SUT,^44^ a mixture of 50% w/w sucrose, 25% w/w urea, 10% w/w 2,2’,2”,-nitrilotriethanol, and 0.1% v/v Triton X-100 (designated ScaleCUBIC-2) for SeeNet^41^ and CUBIC^46^; the same components as the delipidation solution for Scale^43^ and FOCM^45^; CUBIC-RA for CUBIC-L;^86,87^ and the Puclear RI-matching solution^29^ for CUBIC-LH.

### Whole-body clearing

#### CUBIC-L

##### Perfusion and Fixation

We transcardially perfused mice with 40 ml of 0.1 M PBS to flush blood and then fixed them with 150 ml of cold 4% PFA in 0.1 M PBS at a rate of 15 ml/min. We followed this with a perfusion of 20 ml of 0.1 M PBS at a rate of 10 ml/min to remove residual PFA.

##### Delipidation

We transcardially perfused the fixed mice with 100 ml of 1/2-diluted CUBIC-L in dH₂O at a rate of 10 ml/min. Subsequently, we incubated the mice with 200 ml of 1/2-CUBIC-L at 37 °C while shaking at 60 r.p.m. for 6 hours. We continued incubation with CUBIC-L under the same conditions for 6 days, refreshing the solution every 2 days.^47^

##### RI Matching

We washed the cleared samples three times with 0.1 M PBS for 8 hours each to remove residual clearing reagents. The samples were then incubated in 1/2-CUBIC-RA for 1 day, followed by an additional 2 days in CUBIC-RA, with gentle shaking at room temperature. The 1/2-CUBIC-RA consists of an equal mixture of dH₂O and CUBIC-RA. CUBIC-RA contains 45% w/v antipyrine and 30% w/v N-methylnicotinamide, with the pH adjusted to 8.5 using N-butyldiethanolamine, resulting in a final RI of 1.52.^87^ Finally, we incubated the samples in CUBIC-RA or Puclear RI-matching solution for 3 days at room temperature.

#### ARCHmap

The perfusion and fixation processes followed the protocols outlined in the Perfusion and Postfixation section.

##### Decalcification

We intracardially flushed the postfixed mice with 0.1 M PBS for 1 hour. We then subjected them to cardiovascular circulation with 11.5% and 23% w/v ethylenediaminetetraacetic acid disodium salt (EDTA-Na₂, equivalent to 10% and 20% w/v ethylenediaminetetraacetic acid) solutions diluted in dH₂O at 37 °C for 2 days each. We adjusted the pH of the solution to 8.0 using sodium hydroxide and refreshed the decalcification solution daily. After decalcification, we flushed the samples with 0.1 M PBS three times for 3 hours each to remove residual EDTA-Na₂.

##### Delipidation

We developed and integrated whole-body clearing protocols based on CUBIC-L.^39,88,89^ We intracardially circulated whole-body samples with CUBIC-LH at 37 °C for 6 days, refreshing the solution daily for the first 2 days and every 2 days thereafter. After clearing, we circulated the samples with 0.1 M PBS at 37 °C three times for 8 hours each to remove residual clearing reagents.

##### Embedding

We freshly prepared the embedding monomer solution, achieving a final concentration of 4% HMS-iohexol solution and 25% w/v BSA. We mixed equal volumes of 8% HMS-iohexol solution and 50% w/v BSA solution to create the HMS and BSA mixture. The 8% HMS-iohexol solution comprises 67% w/v iohexol, 4% w/v PFA, 8% w/v acrylamide, 0.1% w/v N,N′-methylenebisacrylamide, and 0.5% w/v VA-044 in 0.1 M PBS. To embed the cleared whole-body samples, we injected the abdomen with the embedding solution. We subsequently immersed the entire sample in an additional 250 ml of the embedding solution at 4 °C for 2 days with gentle shaking. Afterward, we transferred the whole-body samples to a mold coated with hydrophobic material and immersed them in 100 ml of fresh embedding solution. Finally, we vacuumized the samples on ice for 20 minutes and polymerized them at 37 °C for 4 hours.

To compare embedding conditions, we mixed different concentrations of HMS solution and BSA solution. The conditions included H0B0 (no embedding), H2B10 (2% HMS-iohexol and 10% w/v BSA), H2B25 (2% HMS-iohexol and 25% w/v BSA), H4B10 (4% HMS-iohexol and 10% w/v BSA), and H4B25 (4% HMS-iohexol and 25% w/v BSA).

##### RI Matching

We utilized the previously reported Puclear RI-matching solution for this experiment.^29^ We fully immersed the embedded samples in the Puclear RI-matching solution at room temperature, gently shaking for at least 10 days before imaging. We refreshed the solution 3 days prior to imaging. The samples could be stored in the Puclear RI-matching solution for over two months in the dark without significant fluorescence quenching.

### Whole-body immunohistochemistry

To perform immunostaining of the sympathetic nerve, we inserted catheters into both the left ventricle and the upper abdomen of the cleared mouse. We then perfused the mouse with a pretreating solution (0.2% v/v Triton X-100, 20% v/v DMSO, 0.3 M glycine, and 0.005% w/v NaN_3_ in 0.1 M PBS) at 37 °C for 1 day. After this, we blocked the samples in a blocking solution (0.2% v/v Triton X-100, 10% v/v DMSO, 0.005% w/v NaN_3_, and 3% v/v donkey serum in 0.1 M PBS) at 37 °C for an additional day. We washed the samples three times with PTwH (0.2% v/v Tween-20, 10 mg/ml heparin, and 0.005% w/v NaN_3_ in 0.1 M PBS), each wash lasting 1 hour. For primary antibody staining, we incubated the samples with 200 ml of rabbit anti-tyrosine hydroxylase (TH) antibody (1:500 dilution) in an incubation solution (5% v/v DMSO, 0.005% w/v NaN_3_, and 2% v/v donkey serum in PTwH) at 37 °C for 7 days. To facilitate the outflow of the incubation solution, we made a small puncture in the lower abdomen. After removing the primary antibody solution, we washed the samples three times with PTwH for 8 hours each at 37 °C. Next, we circulated the samples with the incubation solution containing a secondary antibody (1:500 dilution) for 5 days. Following this step, we washed the samples three times with the washing solution for 8 hours each to remove any residual secondary antibody before the embedding process. All labeling and washing procedures were conducted via active perfusion through the catheters.

To verify the efficiency and specificity of the whole-body immunohistochemistry, we immunostained a DBH-Cre;Ai6 mouse, which expressed ZsGreen in peripheral sympathetic postganglia and fibers, using the same procedures. We examined the colocalization of TH^+^ and ZsGreen^+^ signals in sympathetic ganglia and fibers throughout the entire body.

### Blockface-VISoR system

We developed a custom-built fluorescence microscope system for the automatic imaging and sectioning of whole mouse specimens. This system incorporates an imaging module that utilizes our previously developed VISoR technology.^11,29^ The setup features four lasers operating at 405, 488, 561, and 640 nm (OBIS, Coherent) for fluorescence excitation. We employed a galvo mirror (GVS011, Thorlabs) to generate scanning excitation through an illumination objective (UMPLFLN10XW, NA 0.3, Olympus). We collected fluorescent signals using another objective (UMPLFLN10XW, NA 0.3 or UMPLFLN20XW, NA 0.5, Olympus) positioned perpendicularly to the illumination objective. Both objectives were oriented at a 45° angle relative to the samples. To achieve demagnification and enhance the field of view, we used a 0.63 adapter (TV0.63, Olympus). A scientific CMOS camera (Flash 4.0 v.3, Hamamatsu) captured images continuously at a rate of 400 M pixels per second while the sample was translated at a constant speed using an x-stage (LTS150/M, Thorlabs). We achieved y-translation with a long-range translation stage (LTS300/M, Thorlabs) in a stepwise manner. Additionally, the system includes a sectioning module that cuts samples at a specified thickness at the end of each imaging cycle. This module features a custom-designed vibratome equipped with a tungsten alloy blade driven by an eccentric shaft. The sample apparatus (Figure 1E) comprises a chamber containing RI-matching reagents for imaging, a stainless tube dispenser with an inner diameter of 60 mm, and a piston for elevating the sample. During sectioning, the sample apparatus translates against the blade, utilizing the same y-translation stage (LTS300/M, Thorlabs) to move the sample apparatus back and forth from the imaging module.

### Whole mouse imaging

We secured the cleared whole-body samples onto the piston within the tube dispenser using adhesive. We filled the gap between the sample and the tube dispenser with 4% w/v melted type I-B agarose in RIMS.^39^ During each imaging cycle, we set the x-stage speed in the VISoR imaging module to 0.7 mm/s, while the frame rate of the CMOS camera was set to 200 fps. This setup allowed us to image a sample column measuring approximately 2100 μm in width and 600 μm in depth at a voxel resolution of 1 × 1 × 2.5 μm^3^. Typically, the raw data for an entire cross-section of the mouse body comprised 5 to 10 imaging columns. After capturing the entire cross-section image, the system autonomously moved the sample to the vibratome to trim off the 400-μm top layer. The mouse sample then returned to the VISoR imaging module for the next imaging cycle. This process created an approximately 200-μm-thick overlap between two contiguous imaged sections. We repeated this cyclic process until we completely imaged the whole mouse sample. In total, we successfully imaged 52 adult mice, 11 of which were chosen for further analysis and visualization in this work.

### Tissue size measurement

We assessed linear expansion or shrinkage of the sample using bright-field images. We outlined the borders of the slices with the “polygon selection” function in ImageJ/Fiji. To determine the linear size change, we calculated the ratio of the square root of the area after reagent treatment to the area before reagent treatment.

### Fluorescence intensity

We resampled Z-projections of 30-μm-thick 3D imaging data blocks. We measured the mean signal and background grayscale values from 500 × 500-μm^2^ areas in multiple cortical regions, including the motor, agranular insular, somatosensory, visceral, auditory, and posterior parietal association cortices. We identified soma signals and background using the threshold function in ImageJ. We defined fluorescence intensity as *ΔF = F_Signal_ - F_Background_,* where “*F_Signal_*” and “*F_Background_*” represent the average fluorescence intensities of the signals and background, respectively. We calculated the normalized fluorescence intensity ratio as the *ΔF* of one group relative to that of the PBS group.

### Whole-body reconstruction

The whole mouse body comprised approximately 200 3D cross-sections, with each cross-section stitched together from 5 to 10 raw imaging columns by minimizing 3D translation errors between adjacent columns. We developed a coarse-to-fine registration method for reconstructing the cross-sections, leveraging the overlapped data acquired from this system. The coarse image registration identifies similar surfaces in adjacent sections by matching overlapping voxels. We selected pairs of 250 × 250 × 40 voxel imaging blocks from the overlapping volume of two contiguous imaged sections at horizontal and vertical intervals of 500 pixels. We aligned the blocks from the lower section to their corresponding blocks in the upper section to determine the displacement parameters based on normalized cross-correlation (NCC). A surface deformation field was generated from the matrix of displacement parameters using B-spline interpolation. Next, we joined the two adjacent sections through fine registration of the two surfaces defined within the sections. One surface was a flat surface within the upper section, located at a relative height of 40 μm from the bottom, while the other was the deformed upper surface of the lower section, extracted according to the deformation field at a relative height of 440 μm from the bottom of the section. We selected offsets of 40 μm and 440 μm to avoid defects in the raw data associated with the true sample surfaces. We generated a new deformation field to achieve precise registration between two surface images using a B-spline-transformation implemented via SimpleElastix.^83^ We globally optimized this deformation field across all sections using the stochastic gradient descent algorithm, minimizing the average deformation of the surface image and the average displacement between the upper and lower surfaces of each section. Finally, we employed linear interpolation to extend the refined multiple deformation fields into a comprehensive 3D deformation field. This 3D deformation field was mapped onto the entire dataset, enabling complete 3D reconstruction of the whole mouse body.

We converted the whole-body images or their regions of interest (ROIs) to the Imaris file format for subsequent 3D visualization, rendering, contouring, and animation using Imaris software. The Imaris surface tool facilitated the segmentation of specific tissue types by manually delineating the borders. We then automatically generated the surfaces of the masked volumes. Representative 2D images were created using ImageJ, and Lychnis was employed to animate the whole-body and whole-head images.

### Thoracic skeleton alignment

We achieved alignment of the thoracic skeleton between the target and subject mouse through an iterative approach based on smoothing thin plate spline (STPS).^90^ This method involved: (1) manually marking approximately 200 groups of points at corresponding anatomical locations on the thoracic skeletons of the two mice using the Vaa3D tool;^82^ (2) utilizing STPS to obtain the deformation field between the two groups of points to warp the thoracic skeleton of the subject mouse; and (3) iteratively replacing the original thoracic skeleton of the subject mouse with the warped version until we achieved alignment. To prevent distorted deformation fields during the iteration process, we converted the points back to the original image before each warping step.

### Axonal tracing

We conducted single-neuron tracing using the Lychnis software, as previously reported.^29^ We semi-automatically annotated the entire morphological structures of the nerves at a resolution of 1 μm, enabling the construction of the complete projection route of each individual neuron. All traced neurons were displayed on the surface of one mouse after aligning the thoracic skeleton. We performed image rendering using Imaris. A downsampled reconstructed whole-body image was generated at voxel resolutions of either 10 × 10 × 10 μm^3^ or 4 × 4 × 4 μm^3^, serving as panoramas of high-resolution ROIs for further analysis.

## Notes

https://mesoanatomy.org/mesomouse/meso-example

### Competing Interest Statement

The authors have declared no competing interest.

