## Supplementary figures and legends for "High-speed mapping of whole-mouse peripheral nerves at subcellular resolution"

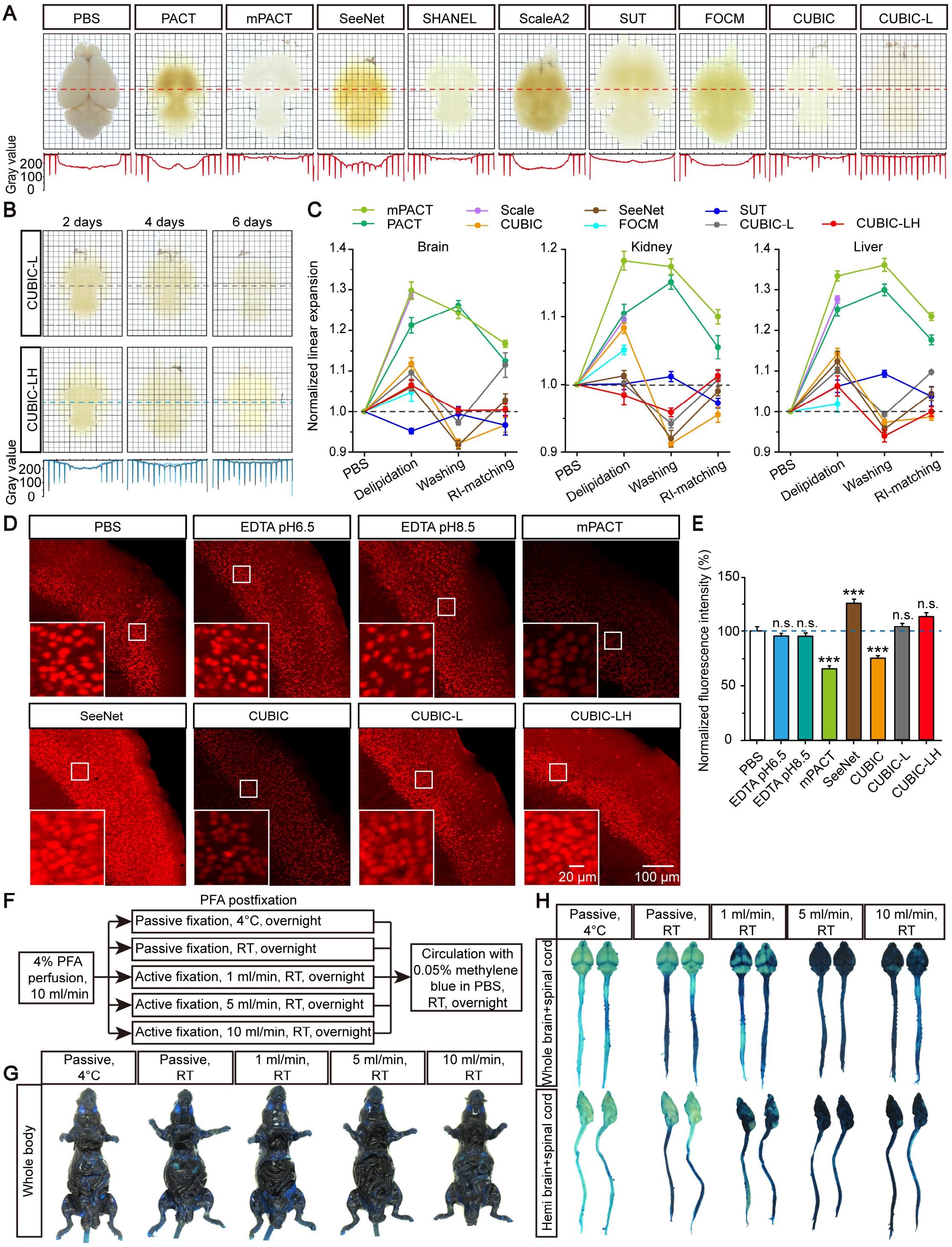


**Figure S1. High-efficiency CUBIC-LH clearing**

(A) Comparison of different aqueous-based whole-brain clearing methods. Grayscale values are measured along the dashed line.

(B) Comparison of the progressive efficiency of whole-brain clearing via CUBIC-L and CUBIC-LH.

(C) Quantitative analysis of the linear change in tissue slices via different aqueous-based clearing reagents following sequential treatments, including delipidation, washing, and RI-matching. n = 4 to 8 slices from 3 mice. Data are presented as mean ± SEM.

(D) Representative fluorescent images from CaMKIIα-Cre;Ai14 brain slices incubated with PBS, decalcification, and delipidation solution for 4 days. Insert panels are magnified tdTomato+ soma in cortical regions. Imaging conditions and contrasts remained the same among different groups.

(E) Quantitative analysis of endogenous fluorescence intensity. n = 52, 63, 41, 77, 80, 76, 80, and 68 regions with a size of 500 × 500 μm2 from 3 mice for the PBS, EDTA pH 6.5, EDTA pH 8.5, mPACT, SeeNet, CUBIC, CUBIC-L, and CUBIC-LH groups, respectively. One-way ANOVA followed by Bonferroni post hoc test; n.s., no significance; ***$P < 0.001$.

(F) Schematic diagram illustrating postfixation and permeation testing experimental procedures.

(G) Comparison of permeation to viscera via cardiovascular circulation of methylene blue.

(H) Comparison of permeation to the brain and spinal cord extracted from whole bodies in (G).

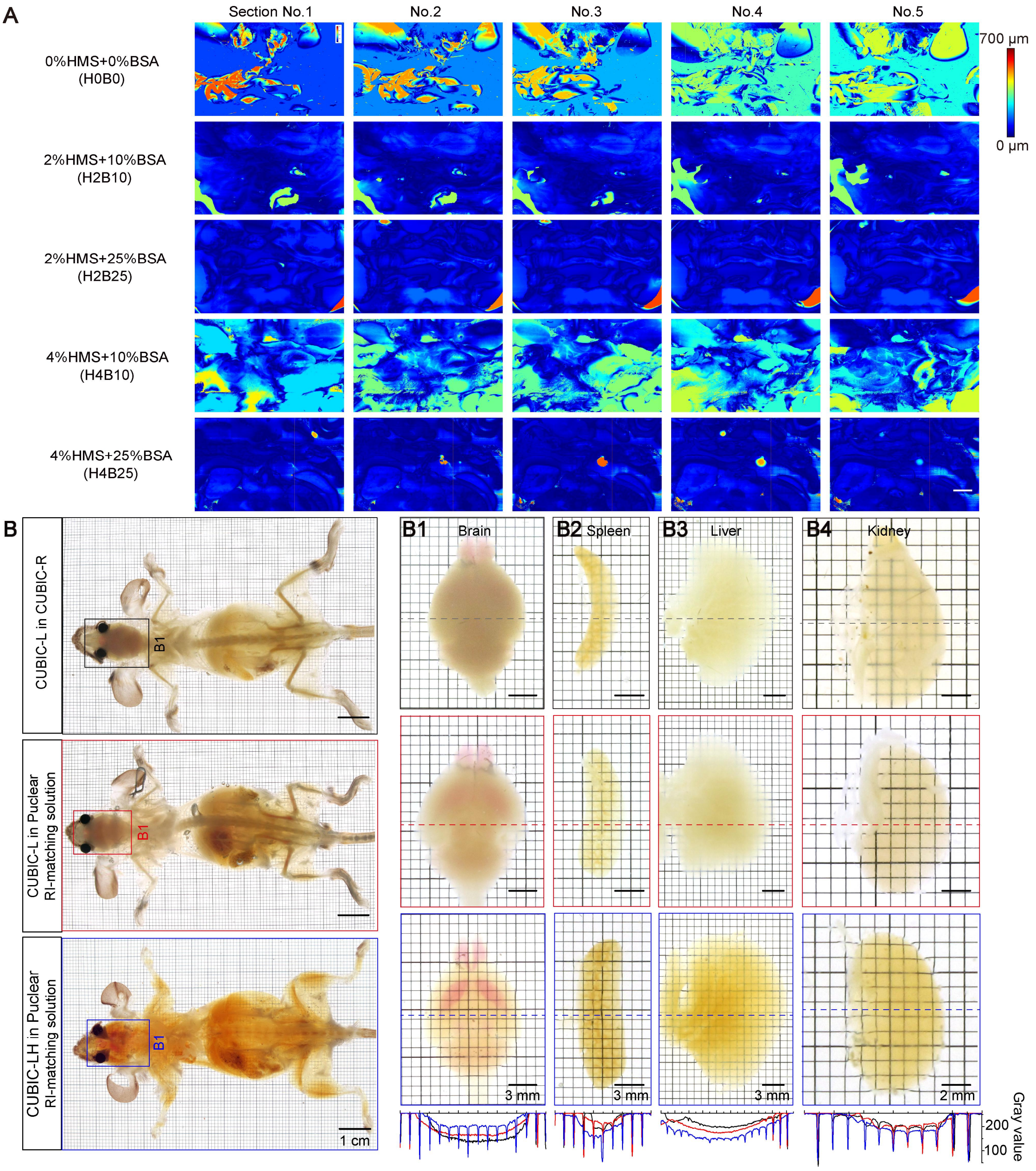


**Figure S2. Whole-body embedding and clearing tests**

(A) Surface topograph of sections processed under various embedding conditions. Color bar indicates different values of depth between each surface pixel and their average values.

(B, B1-B4) Comparison of different whole-body clearing methods. Brain (B1), spleen (B2), liver (B3), and kidney (B4) were extracted from the cleared whole-mouse samples in (B) for transparency comparisons. The whole-body sample and extracted brain cleared by CUBIC-LH in the Puclear RI-matching solution in lower panels are the of those in Figure 1B, which is repeatedly presented for a clearer comparison with other whole-body clearing techniques.

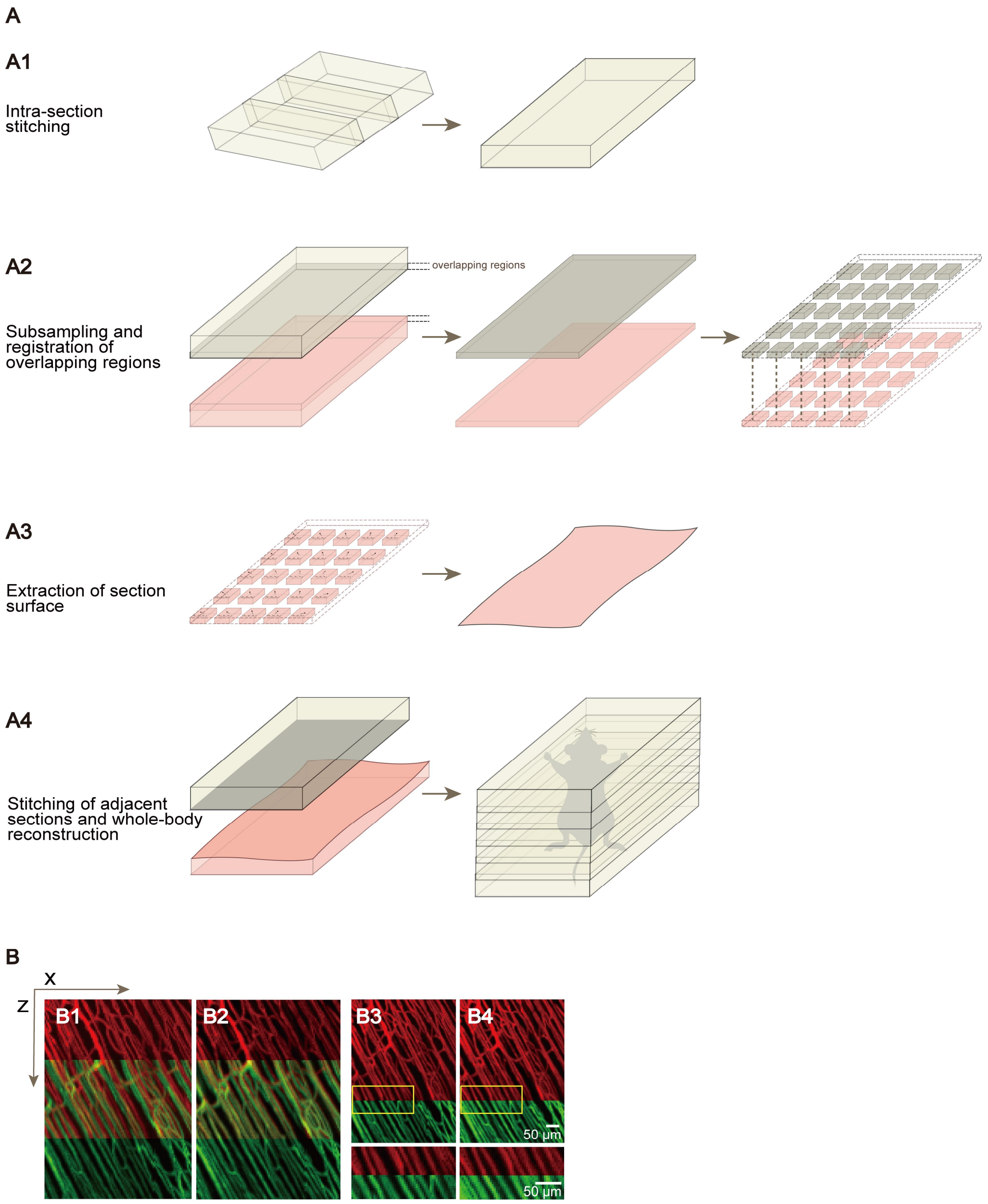


**Figure S3. 3D reconstruction of whole-body data**

(A, A1-A4) 3D reconstruction pipeline for whole mouse bodies. (A1) Intra-section stitching between two imaging columns. (A2) Sub sampling and displacement calculation based on normalized cross-correlation between 2 blocks in the overlapping region from adjacent sections. (A3) Determination of the surface deformation from the displacement of the lower section. (A4) Stitching of adjacent sections and reconstruction of the whole body.

(B) Maximum intensity projection of vessel fluorescence of 200-μm-thick virtual stacks from 2 contiguous imaging sections (red and green, respectively) for comparison of reconstruction qualities using different methods, including natural coordinates (B1, B3) and the customed reconstruction method (B2, B4). In (B1) and (B2), overlapped regions from red and green sections are both presented to directly compare co-registration of fine structures. In (B3) and (B4), the two contiguous sections are stitched to illustrate stitching accuracy.

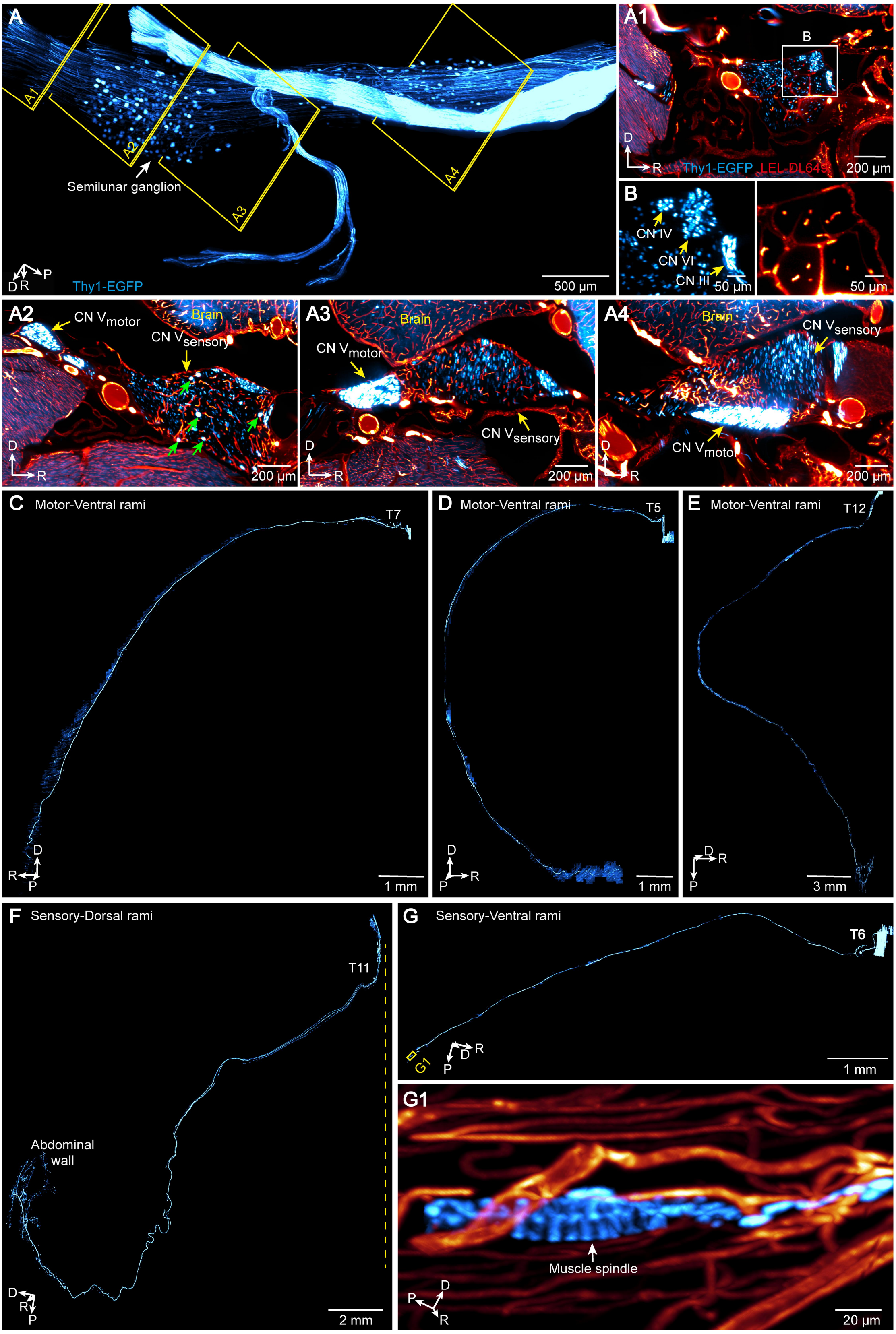


**Figure S4. Visualization of the trigeminal and spinal nerves**

(A) 3D view of the semilunar ganglia, roots, and divisions of the left trigeminal nerve (CN V) in a Thy1-EGFP mouse.

(A1-A4, B) Maximum intensity projections of 20-μm z-stacks boxed in (A), illustrating the spatial distribution of the motor and sensory fibers of the CN V. Red hot (black-red-yellow-light yellow spectrum), LEL-DL649; cyan hot (black-blue-cyan-white spectrum), Thy1-EGFP. Green arrows indicate neuronal somas in the semilunar ganglia. (B) Magnified roots of the oculomotor (CN III), trochlear (CN IV), and abducent (CN VI) nerves coursing within the cavernous sinus in (A1).

(C-E) Representative anatomical morphology of individual spinal motor neurons in the ventral rami. (C) and (E) are viewed in a horizontal mirror-reversed perspective.

(F) Representative anatomical morphology of an individual spinal sensory neuron in the dorsal rami innervating the abdominal skin. Yellow dashed lines indicate the midspinal line.

(G, G1) Representative anatomical morphology of an individual muscle spindle neuron. (G1) Magnified muscle spindle ending from another perspective.

Orientation of images: R: right; P: posterior; D: dorsal.

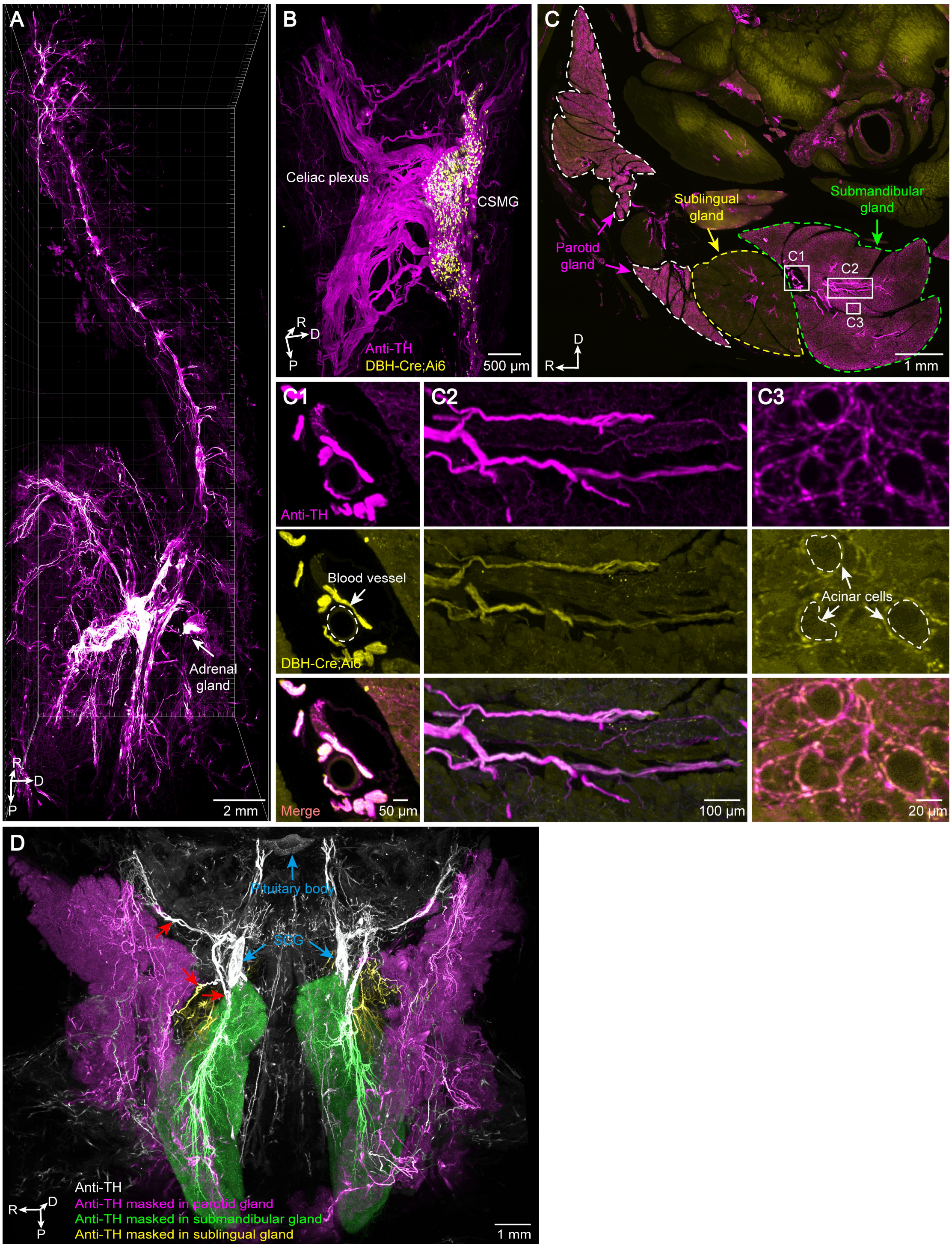


**Figure S5. Uniform whole-body immunostaining of sympathetic nerves**

(A) Lateral views of the segmented sympathetic chain and plexuses.

(B) 3D view of the TH+ celiac-superior mesenteric ganglia (CSMG) and celiac plexus in a DBH-Cre;Ai6 mouse.

(C) Representative images of the maximum intensity projections of a 100-μm coronal section demonstrating distinguished intensity of sympathetic nerves in the submandibular, sublingual, and parotid glands.

(C1-C3) Magnification of colocalized TH+ signals with endogenous ZsGreen (DBH-Cre; Ai6) in perivascular sympathetic nerves (C1), fiber tracts (C2), and endings surrounding acinar cells (C3) in (C).

(D) 3D view of sympathetic innervations to the submandibular, sublingual, and parotid glands, segmented in green, yellow, and purple pseudocolors, respectively. SCG: superior cervical ganglion (lower blue arrows). Red arrows indicate the nerve branches from the SCG to the salivary glands.

Orientation of images: R: right; P: posterior; D: dorsal.

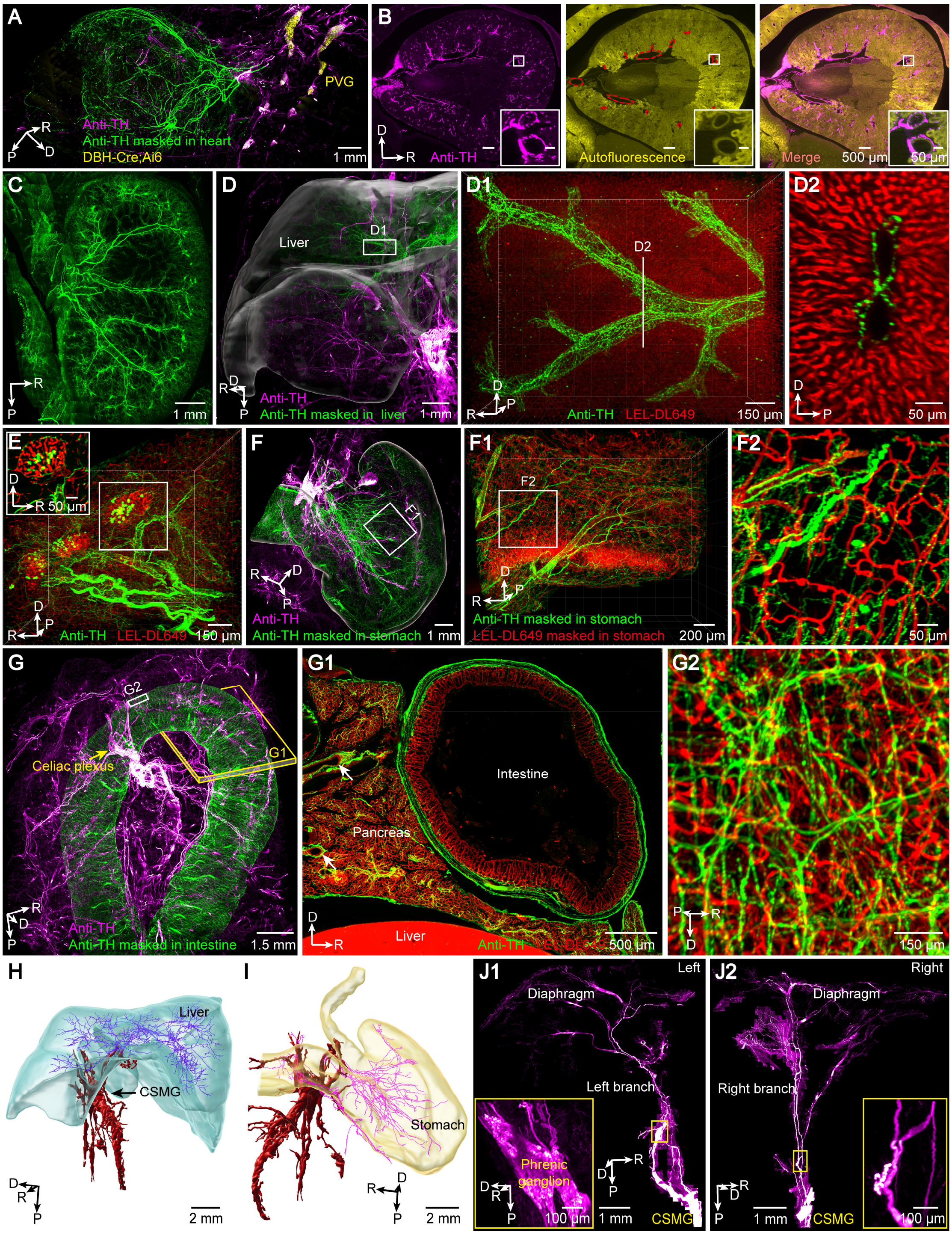


**Figure S6. Sympathetic innervations to visceral organs**

(A) 3D view of sympathetic innervations from the thoracic paravertebral ganglia (PVG) to the heart, segmented in green.

(B1-B3) Representative images of the maximum intensity projections of 10-μm z-stacks showing the perivascular sympathetic plexuses in the kidney. Renal vessels are indicated by red dashed contours.

(C) Maximum intensity projection of 1-mm horizontal sections showing the sympathetic innervations to the kidney.

(D) 3D view of sympathetic innervations to the right lobe of the liver, segmented in green.

(D1) Magnified perivascular sympathetic networks surrounding hepatic blood vessels in (D), displayed from a top perspective.

(D2) Maximum intensity projection of 5-μm-thick stacks indicated in (D1) showing the cross section of perivascular sympathetic nerves.

(E) 3D view of sympathetic nerves and vasculature in the endocrine and exocrine pancreas. Magnified insert indicates the maximum intensity projection of 10-μm-thick stacks showing sympathetic nerves and TH+ endocrine cells in an islet of Langerhans.

(F) 3D view of sympathetic innervations to the stomach, segmented in green.

(F1, F2) Magnified surface view of sympathetic networks and vasculature (zoomed-in view in F2) in the gastric wall in (F).

(G) 3D view of sympathetic innervations to the intestine, segmented in green.

(G1) Maximum intensity projection of a 100-μm coronal section in (G), revealing the distribution of sympathetic nerves in the cross section of the intestine and pancreas.

(G2) Magnified surface view of sympathetic nerves and vasculature in the intestinal wall in (G).

(H-I) Axonal tracing of sympathetic innervations to the liver (H) and stomach (I) from the CSMG.

(J1-J2) 3D view of sympathetic innervations to the diaphragm of a mouse. The left branch is from the left phrenic ganglion (J1), and the right branch is from the CSMG (J2). Magnified inserts illustrate the phrenic ganglion in (J1) and a lack of ganglia in (J2).

Orientation of images: R: right; P: posterior; D: dorsal.

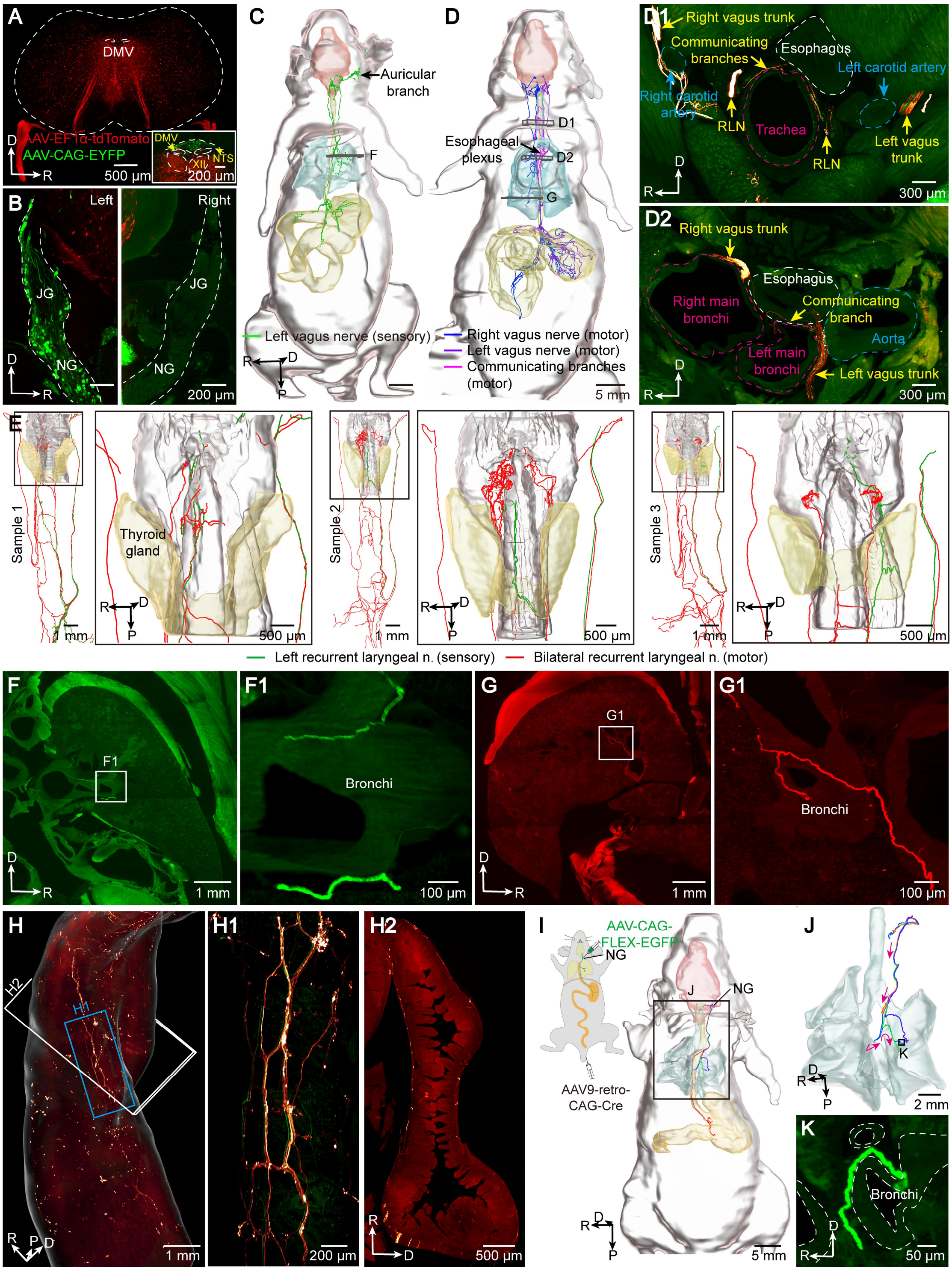


**Figure S7. Vagal motor and sensory innervations in the whole body**

(A) Representative images of the maximum intensity projections of 400-μm z-stacks showing the visceral motor fibers of the brainstem vagus nerve. Insert images are the maximum intensity projections of 10-μm z-stacks, showing AAV-EF1α-tdTomato injection in the DMV with leakage to the adjacent hypoglossal nuclei (XII).

(B) Representative images of the maximum intensity projections of 10-μm z-stacks showing AAV-CAG-EYFP infection of the vagal ganglia.

(C) Whole-body tracing of the visceral and somatic sensory fibers of the vagus nerve to the auris, larynx, lung, stomach, and intestines.

(D) Whole-body tracing of general visceral motor fibers of the vagus nerve in another mouse.

(D1-D2) Maximum intensity projections of 800-μm and 500-μm coronal sections in (D), showing communicating branches between the bilateral recurrent laryngeal nerves and vagal trunks contributing to the tracheoesophageal (D1) and pulmonary plexus (D2), respectively.

(E) Fiber tracing of vagal sensory and motor innervations to the larynx in three samples.

(F-G, F1-G1) Maximum intensity projection of 200-μm z-stacks showing the visceral sensory fibers (F, and zoomed-in view in F1) and visceral motor fibers (G, and zoomed-in view in G1) of the vagus nerve surrounding the bronchi in (C) and (D), respectively.

(H, H1-H2) 3D view (H, zoomed-in view in H1) of vagal fibers in the duodenum. (H2) Maximum intensity projection of 100-μm z-stacks illustrating the distribution of vagal fibers in the cross section. White surface, duodenum.

(I) Single-neuron tracing of six individual vagal sensory neurons across the whole body. Schematic diagram shows the sparse viral labeling procedures of the visceral sensory fibers of the vagus nerve via AAV9-retro-CAG-Cre injection into the tail vein and AAV-CAG-FLEX-EGFP injection into the vagal ganglia.

(J) Magnified five bronchi-projecting vagal sensory neurons in (J).

(K) Magnified sensory ending surrounding the bronchi (white dashed lines) of an individual vagus neuron in (J) from another perspective.

Orientation of images: R: right; P: posterior; D: dorsal.

**Supplementary videos**

**Supplementary Video 1.** Blockface-VISoR imaging
**Supplementary Video 2.** Maximum projection of coronal sections from lectin-labeled Thy1-EGFP mouse
**Supplementary Video 3.** Individual neuron tracing of thoracic spinal nerves
**Supplementary Video 4.** Individual neuron tracing of thoracic spinal nerves
**Supplementary Video 5.** Sympathetic nerves in head and body trunk
**Supplementary Video 6.** Sympathetic innervations to organs
**Supplementary Video 7.** Vagal innervations to organs
